# Machine learning on multiple epigenetic features reveals H3K27Ac as a driver of gene expression prediction across patients with glioblastoma

**DOI:** 10.1101/2024.06.25.600585

**Authors:** Yusuke Suita, Hardy Bright, Yuan Pu, Merih Deniz Toruner, Jordan Idehen, Nikos Tapinos, Ritambhara Singh

## Abstract

Cancer cells show remarkable plasticity and can switch lineages in response to the tumor microenvironment. Cellular plasticity drives invasiveness and metastasis and helps cancer cells to evade therapy by developing resistance to radiation and cytotoxic chemotherapy. Increased understanding of cell fate determination through epigenetic reprogramming is critical to discover how cancer cells achieve transcriptomic and phenotypic plasticity.

Glioblastoma is a perfect example of cancer evolution where cells retain an inherent level of plasticity through activation or maintenance of progenitor developmental programs. However, the principles governing epigenetic drivers of cellular plasticity in glioblastoma remain poorly understood. Here, using machine learning (ML) we employ cross-patient prediction of transcript expression using a combination of epigenetic features (ATAC-seq, CTCF ChIP-seq, RNAPII ChIP-seq, H3K27Ac ChIP-seq, and RNA-seq) of glioblastoma stem cells (GSCs). We investigate different ML and deep learning (DL) models for this task and build our final pipeline using XGBoost. The model trained on one patient generalizes to another one suggesting that the epigenetic signals governing gene transcription are consistent across patients even if GSCs can be very different. We demonstrate that H3K27Ac is the epigenetic feature providing the most significant contribution to cross-patient prediction of gene expression. In addition, using H3K27Ac signals from patients-derived GSCs, we can predict gene expression of human neural crest stem cells suggesting a shared developmental epigenetic trajectory between subpopulations of these malignant and benign stem cells.

Our cross-patient ML/DL models determine weighted patterns of influence of epigenetic marks on gene expression across patients with glioblastoma and between GSCs and neural crest stem cells. We propose that broader application of this analysis could reshape our view of glioblastoma tumor evolution and inform the design of new epigenetic targeting therapies.

## 1. Author summary

This study aimed to develop a machine learning (ML) pipeline that can be used to investigate the role of epigenetic regulation on gene transcription in glioblastoma stem cells (GSCs). We developed a cross-patient prediction pipeline with multi-epigenomic data of patient-derived GSCs to predict gene expression. Our pipeline includes in-silico perturbation analysis, which examines the impact of different epigenetic regulators including chromatin accessibility (ATAC-seq), distal chromatin looping (CTCF ChIP-seq), histone modifications (H3K27Ac ChIP-seq), and active transcription (RNAPII ChIP-seq), on gene transcription across patients. Our in-silico perturbation analysis inferred that the various epigenetic modulators are essential for regulating gene expression, with a higher weight on H3K27Ac, across patients. Collectively, we developed a cross-patient prediction pipeline that can be used to unravel the multi-epigenetic-driven mechanism of gene expression and propose potential drivers of cellular plasticity in GSCs.

## 2. Introduction

Glioblastoma stem cells (GSCs) are characterized by tumor-initiating and self-renewal properties and are known to drive chemoresistance and heterogeneity in glioblastoma (1–5). Until now, there is limited understanding regarding the epigenetic factors that define GSCs’ failure to attenuate their stemness potential in the face of differentiation cues. In addition, the influence of epigenetic mechanisms on GSC phenotypic plasticity is not fully understood. To address these questions, previous studies have examined the contribution of histone modifications, chromatin accessibility, and distal chromatin looping on gene transcription (6–9). Conventional studies employ correlation analysis to look at the linear relationship between gene expression and one epigenetic modulator. For example, it was shown that histone modifications can be predictive for gene expression by looking at the linear relationship between expression and histone modifications (10). In addition, CTCF was shown to play a role in gene regulation by participating in establishment and maintenance of chromatin loops (11). Finally, gene expression of oncogenes is amplified due to increased chromatin accessibility and enhancer activation (12). Although these studies provide mechanistic insights on epigenetics-based modulation, they include poor correlation coefficients between each epigenetic modulator and gene expression suggesting that correlation analysis is incapable of comprehensively interrogating the enormous amount of epigenomic and transcriptomic data.

To address this issue, machine learning (ML) techniques have been applied to epigenomics data such as ATAC-seq, ChIP-seq of histone modification marks, or transcription factors. The majority of these studies have been applied to predict gene expression based on one type of epigenetic modulator, such as histone modifications, as seen in AttentiveChrome and GraphReg (13–15). In the last few years, few studies have also applied ML algorithms to build models that take multiple epigenetic modulators to predict gene expression (16–19). However, the application of machine learning to study the combinatorial effect of multiple epigenetic modulators across patients with cancer has not been performed.

Here, we develop an ML-based prediction model that predicts gene expression levels in patient-derived GSCs using multiple epigenetic regulators (ATAC-seq, RNA polymerase II (RNAPII) ChIP-seq, CTCF ChIP-seq, and H3K27Ac ChIP-seq) at high performance. To examine the contribution of each epigenetic regulator on gene expression, we perform in-silico perturbation analysis and show that all epigenetic regulators contribute to gene expression prediction, with a higher contribution of H3K27Ac ChIP-seq signals, followed by RNAPII ChIP-seq, ATAC-seq, and CTCF ChIP-seq across patients.

Glioblastoma is a perfect example of cancer evolution where cells retain an inherent level of plasticity through activation or maintenance of progenitor developmental programs. To determine the contribution of H3K27Ac and CTCF binding on predicting gene expression between GSCs and neural stem cells we used publicly available ChIP-seq data of H3K27Ac and CTCF of neural crest (NCCs) and neural progenitor cells (NPCs) as test data for our GSC data-trained model and compared the Pearson Correlation Coefficients (PCCs). Our analysis shows that the GSCs and NCCs share patterns of H3K27Ac enhancer landscape influence on their gene expression.

Overall, our ML approach determines weighted patterns of influence of epigenetic marks on gene expression across patients with glioblastoma. Moreover, our approach reveals pattern similarity of H3K27Ac marks across GSCs and NCCs, which provides insights into epigenetic-driven cellular plasticity. Broader application of this analysis could reshape our view of glioblastoma tumor evolution and inform the design of new epigenetic targeting therapies.

## 3. Materials and methods

### 3.1 Datasets and pre-processing

We model and investigate the relationship between epigenetic modulators and gene transcription of patients derived GSCs using machine learning. To achieve this, we used the following four markers to compose our GSC patient datasets: ChIP-sequencing with H3K27Ac (enhancer marker), RNAPII (active transcription marker), and CTCF (distal chromatin looping marker), and ATAC-sequencing (chromatin accessibility) and RNA-sequencing data (Fig 1). This dataset was created for two patients (GSC1 and GSC2). For investigative downstream experiments, we included datasets composed of markers from non-GSC crest and progenitor neural cell data sourced from ENCODE (accession codes: ENCFF056WDN, ENCFF521XJN, ENCFF400MZX, ENCFF123YLB, ENCFF503KKJ, ENCFF655GGB, and ENCFF583OOM)(18,20–26). Both neural cell samples were from the human embryonic stem cells (H9) cell line. The progenitor cell dataset included H3K27Ac ChIP-seq, CTCF ChIP-seq, and DNase-seq (as an analog for ATAC-sequencing). The crest cell dataset included H3K27Ac ChIP-seq and CTCF ChIP-seq. For detailed information on the preparation of these datasets see supplementary section S1.

**Fig 1.**
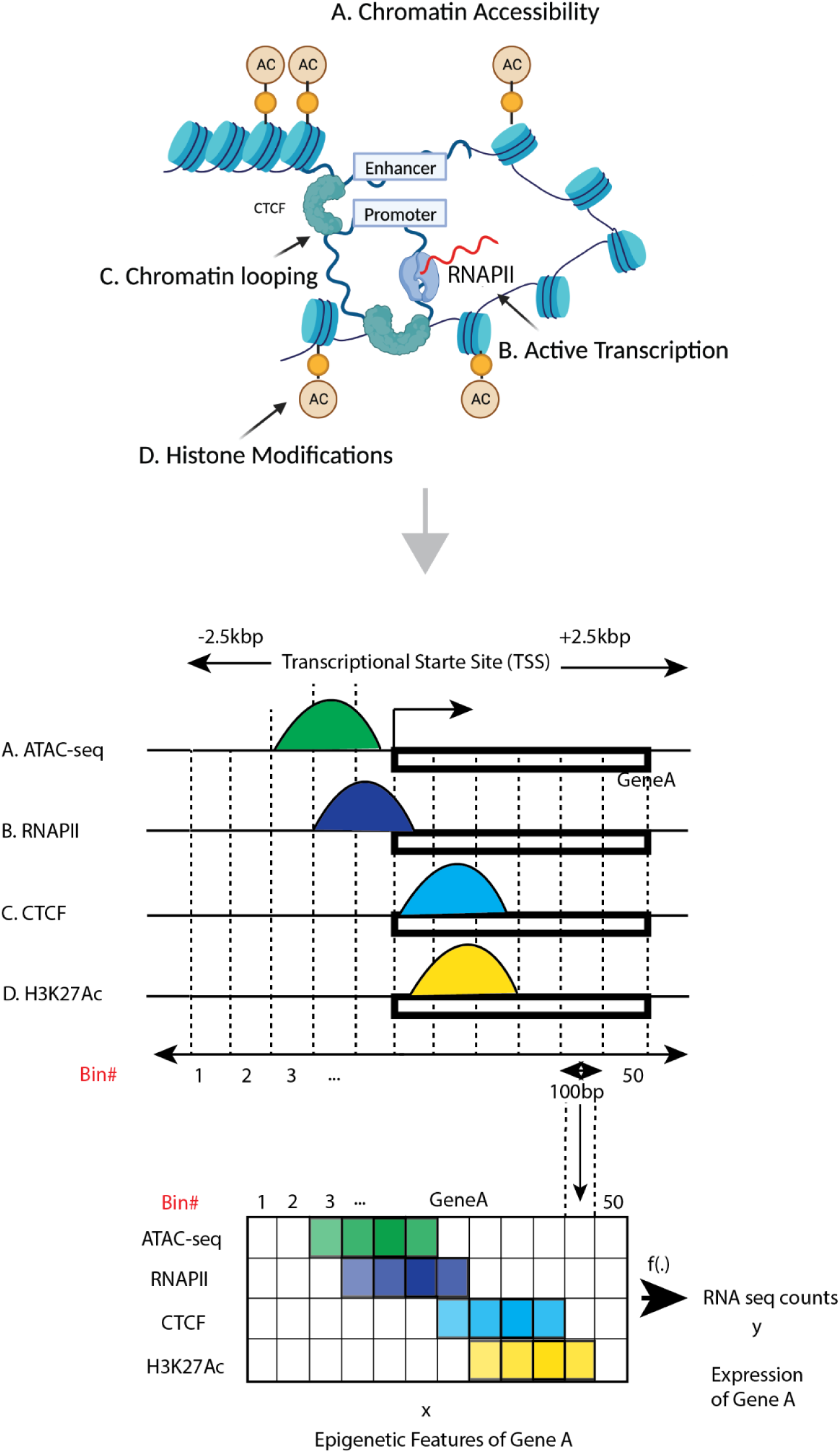
Schematic overview of epigenetics-driven gene transcription and epigenomics sequencing data processing: Gene transcription heavily relies on epigenetics. These epigenetic mechanisms can be categorized into four categories: A. Chromatin accessibility, B. Active Transcription, C. Chromatin looping, D. Histone modifications. Counts of these sequencing +/-2.5 kilo base-pairs (kbp) flanking the TSS region of each gene were measured and divided into 50 bins, with each bin representing 100 base pairs to create a heatmap for the input of the model.

We focused on the +/-2.5 kilo base-pairs (kbp) flanking region of the transcription start site (TSS) for each gene and divided it into 50 bins, with each bin representing 100 base pairs. Further, we created a 50 x 4 matrix with rows representing bins and columns for epigenetic features for each of the 20,015 genes, and each bin contains summarized counts. As a result, we passed 20,015 x 50 x 4 to our model as an input. To prepare the gene expression labels, we summarized counts of the +/-2.5kb flanking the TSS per gene and normalized it using transcripts per million (Fig 2).

**Fig 2.**
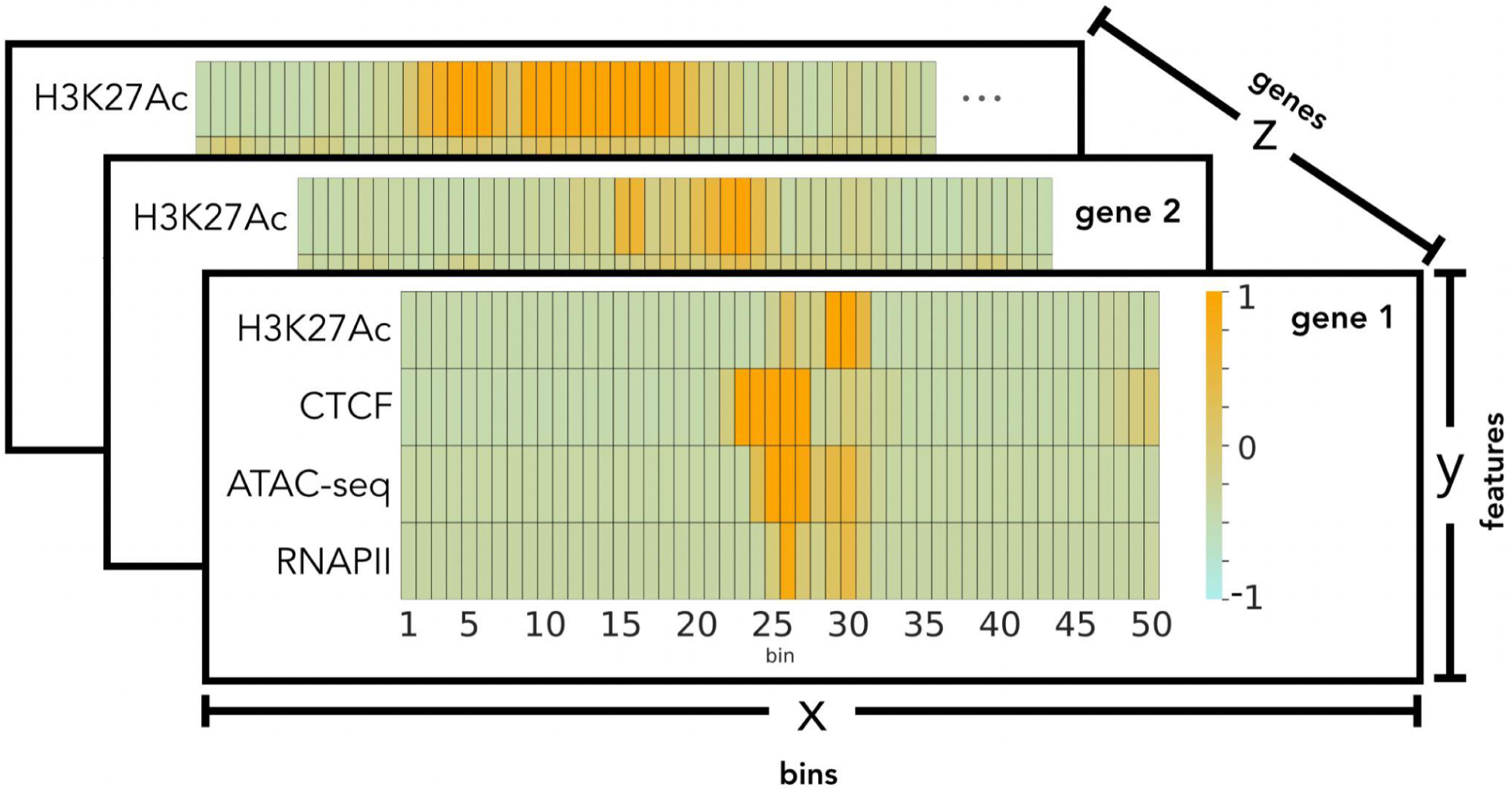
Patient datasets representation after preprocessing. The figure depicts the standardized epigenetic marker values per gene for a patient. It highlights the 3-dimensional arrangement of the datasets prior to model input. Here the “x” axis corresponds to the 50 bins of 100bp counts for each feature. The “y” axis represents each gene’s 4 epigenetic features. The figure’s “z” axis is representative of the gene arrangement in the dataset.

The second preprocessing step included separate standardization and log(2) transformation of the gene expression measurements (27). To account for the variation in count values across different types of sequencing and experimental conditions, we standardized the counts of each sequencing data by using the mean value of the corresponding sequence data. This standardization occurred after the train, validation, and test data-splitting was performed. Here, each of the four epigenetic features were standardized separately within the train, validation, and test sets. We chose to standardize the data at this point, as opposed to before the data splitting, to avoid potential data leakage (28). Each gene had its corresponding target label log(2) transformed with a pseudo count of 1, prior to the data split process. Additional information regarding the model input process is contained in the supplementary section S2 (Fig S1 & S2).

### 3.2 Machine learning modeling

We tested several regression-based cross-patient predictive models to examine the novel application of our specific combination of epigenetic markers to predict gene expression using machine learning. Given the two patients (GSC1 and GSC2), we sought to use machine learning to extract a common pattern seen across patients, as GSCs are highly heterogeneous from one patient to the next. Therefore, we trained each model using a subset of genes from patient 1 followed by testing the same model with patient 2’s dataset (represented as GSC1→GSC2). Our experimental setup also included the inverse operation where the models were trained on patient 2’s dataset and tested using patient 1’s data (represented as GSC2 →GSC1).

To our knowledge this was the first time this specific combination of markers was explored in a machine learning study. In every case, the prediction task was a regression where the models predicted the RNA-seq gene expression value per gene. Cross-patient prediction experiments required two datasets (one for training and validation, and the other for testing), each composed of ChIP-seq, ATAC-seq, and RNA-seq.

To select the best machine learning model, we tested deep learning architectures like a Multi-layered Perceptron (MLP), a Convolutional Neural Network (CNN), a Recurrent Neural Network (RNN) and a Branched Multi-layer Perceptron (Branched MLP). With the branched MLP we combined the genomic sequences from the region as inputs.

Traditional machine learning algorithms included Gradient Boosting Regression (GBR), Support Vector Regression (SVR), and Multiple Linear Regression (MLR) architectures. Detailed information regarding these models is outlined in supplementary section S3.

**Evaluation metrics.** Pearson Correlation Coefficient (PCC) was used as the primary metric for hyperparameter tuning, performance evaluation, and feature perturbation. Spearman Correlation Coefficient (SCC) was calculated concurrently with PCC.

#### 3.2.1 Criteria for model selection

Out of all the models tested, we selected our final machine learning model based on the analysis of each model’s overall Pearson Correlation Coefficient (PCC) metric results.

#### 3.2.2 XGBoost model details

The XGBoost based model exhibited the best average performance of all the cross-patient experiments in this study. The downstream analysis and discussions we share in later sections are drawn from this architecture’s predictions. Since XGBoost takes 2-dimensional input, the dataset features were flattened to 20,0015 x 200 while the target variable (RNA-seq) became a 20,0015 x 1 array. The relative positioning of the bins of each of the four gene features was kept contiguous (Fig 3).

**Fig 3.**
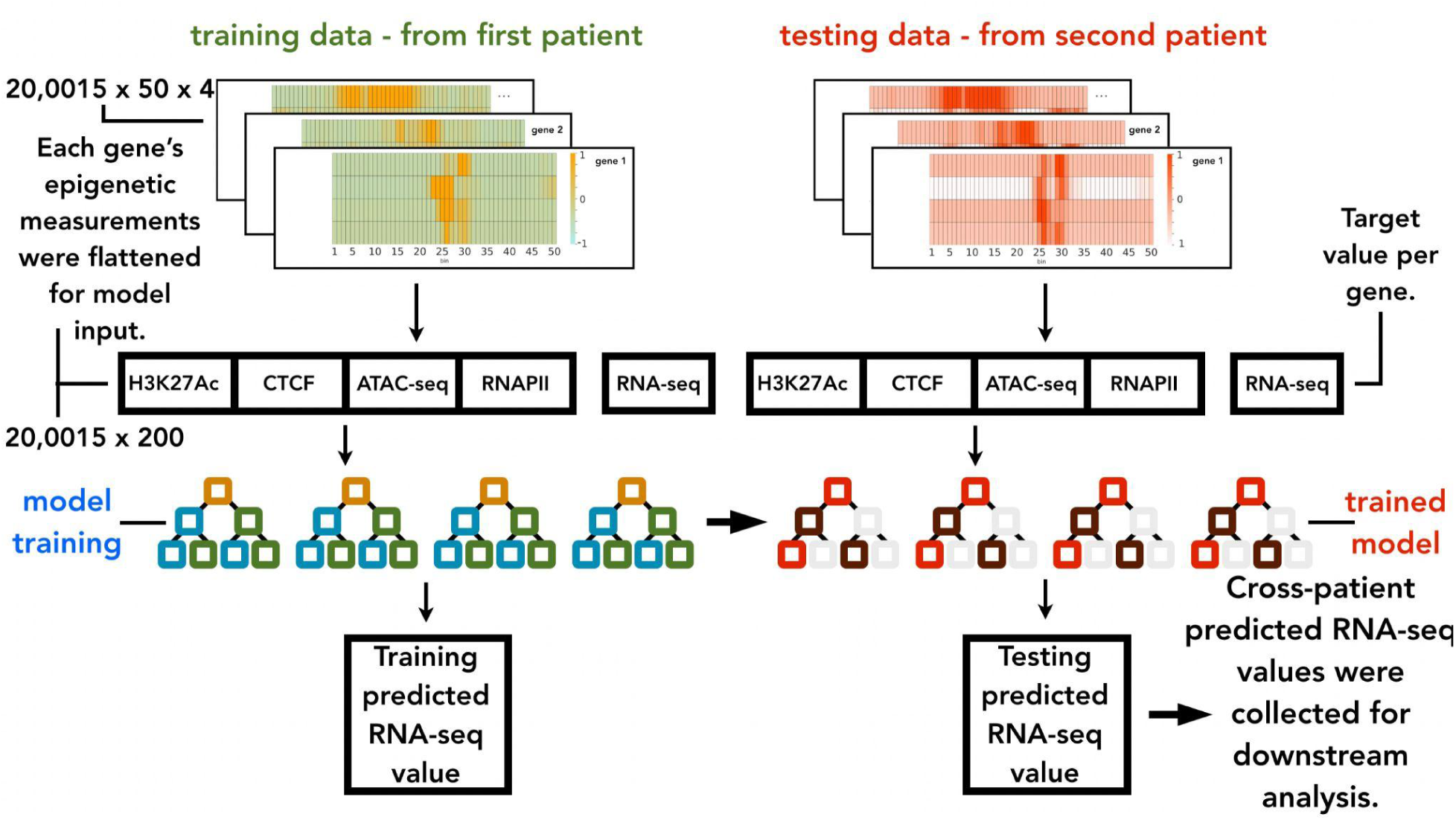
Cross-patient prediction methodology using the model XGBoost architecture. The model input for training and validation was derived from a different patient than the testing dataset. As shown, the matrices were flattened before going into the model where the RNA-seq value was predicted.

**Loss function.** We used mean squared error (MSE) for loss function calculations as follows:

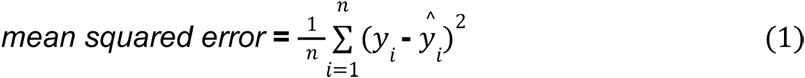

In the calculation *y_i_* represents the actual RNA-seq measurement while 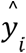 represents the predicted value per gene *i* for the number of genes *n* in the dataset.

### 3.3 Model training, validation and testing

We split the data into training, validation, and test sets to perform hyperparameter tuning. We trained two models (GSC1→GSC2 and GSC2→GSC1). The hyperparameters for the first model were tuned by dividing the GSC1 datasets into training (70%) and validation (30%) sets. All the genes in the GSC2 dataset constituted the test set. Similar setup was used for the GSC2→GSC1 model. A base set of hyperparameters for experimental testing, optimized for higher PCC performance, were chosen for each model. This was done using a grid search (combinatorial products of hyperparameter values into unique sets) optimizing for model performance. Supplementary section S3 (Table S1, S2, S3, S4, S5, S6, S7, and S8) includes additional information on hyperparameter tuning.

We ran each model 10 times (with different random seeds) in the two cross-patient arrangements (GSC1→GSC2 and GSC2→GSC1) for experimental testing. The mean and standard deviation of our metrics are reported for our study’s experimental results (Fig 4A and 4B).

**Fig 4A& 4B.**
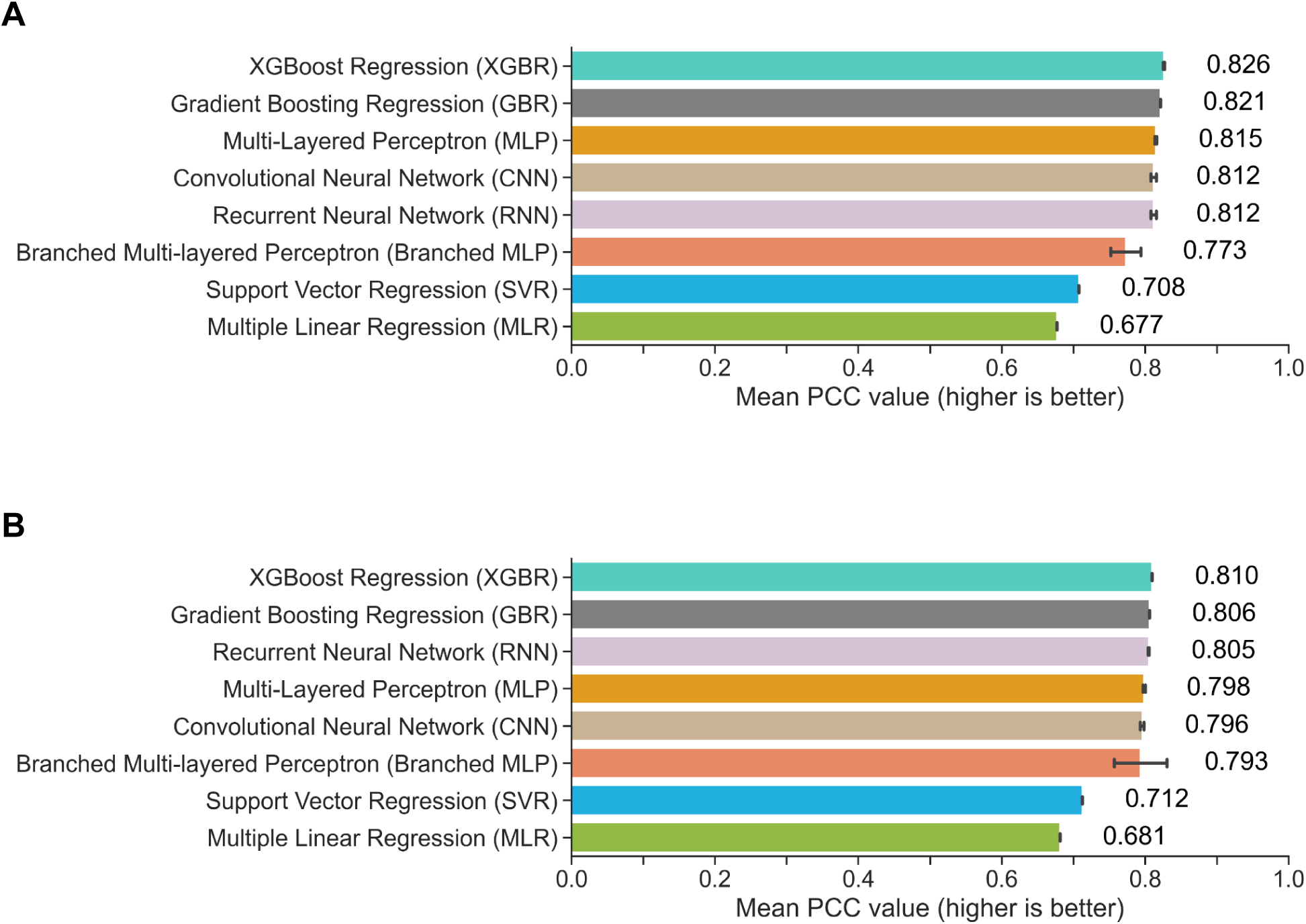
PCC cross-patient regression model results. Our experimental results were compiled as the mean PCC scores over 10 runs of each model. In both graphs, the error bars shown indicate standard deviation of the model results. A) Our cross-patient XGBoost-based regression model performed higher than all other architectures when training with GSC1 and testing with GSC2 (GSC1→GSC2). B) Our XGBoost-based algorithm also out-performed the others when training with GSC2 and testing with GSC1 (GSC2→GSC1).

### 3.4 Perturbation experimental setup and analysis

To study the individual effect of each epigenetic signal on prediction, we perturbed one of each of the datasets four epigenetic features at a time and ran the perturbed input through the trained model. For this set of experiments, each signal perturbation was done for 10 runs while each run was given a different random seed. The perturbation experiments maintained our cross-patient prediction arrangement and were run in both prediction directions (GSC1→GSC2 and GSC2→GSC1).

Perturbation of a particular epigenetic signal was achieved by replacing all of that feature’s values with 0.0 (the mean of the standardization) for all gene’s across the test dataset. Our analysis compared the calculated mean and standard deviation results, for each set of perturbations, to the original model performance and each other. The per-gene prediction results were also recorded for later analysis.

## 4. Results

### 4.1 Machine and deep learning models perform similarly in different cross-patient prediction scenarios

The study found that, when the models were trained with GSC1’s (patient 1) and tested with GSC2’s (patient 2) data, which we referred to as GSC1→GSC2, the highest PCC value was acquired with the XGBoost Regression (XGBR) model at 0.826199 ± 0.000888. Multiple Linear Regression (MLR) held the lowest performance of 0.676872 ± 0.0; a 0.149327 difference in PCC. The sub-par performance of MLR indicated that the relationships among the epigenetic markers and their relationship with gene expression was non-linear. This characteristic of the data reinforced the need to experiment with many different algorithms of higher complexity to optimize our study’s application of machine learning. Interestingly, given the non-linearity of the data, five of our models successfully found patterns leading to RNA-seq values using completely different architectures. In fact, this sub-group (XGBR, GBR, MLP, CNN, and RNN) performed within 0.014497 of each other. This indicated that the results were to some degree agnostic to the model used and more so dependent on the epigenetic features of the dataset.

For the following set of experiments, we trained all of our models with GSC2’s (patient 2) dataset and tested with GSC1’s (patient 1). We referred to this as the GSC2→GSC1 prediction direction. The highest level of performance we achieved with this set of experiments was a PCC score of 0.809537 ± 0.000377, which as it turns out was again from XGBR. Comparing this prediction direction to the previous one we found that the general trends in metrics were similar. The separation between XGBR and MLR was 0.12818 for this prediction direction. Meanwhile, the difference among the first and sixth models was 0.016044. These two findings indicate the small differences in PCC among our higher performing models for this prediction direction as well.

Our data suggest that across both prediction directions, XGBR emerged as the model with the highest test set PCC measurements. Additionally, the strong PCC values for most of our models including XGBR contrasted by the consistently lower than average results from our MLR, underscored the non-linearity of our GSC datasets. Therefore, we used the XGBR model for our following perturbation experiments and downstream analysis. Finally, the strong PCC values highlighted the success of our cross-patient prediction approach to generalize the epigenetic input patterns connected to the gene expression from one patient to another. Our methodology of experimenting with both the GSC1→GSC2 and GSC2→GSC1 prediction directions successfully found high performance and consistency across our metrics.

Our study’s corresponding SCC metric results are detailed in the supplementary section S5 (Fig S4A & S4B). Script time considerations are detailed in the supplementary section S6 (Table S9).

### 4.2 Perturbation Results: All the combined epigenetic markers (epigenetic modification, chromatin accessibility, and histone modifications) contribute to gene expression with a higher weight on H3K27Ac signals

To evaluate the contribution of each epigenetic marker to predicting gene expression, we conducted perturbation analyses on each marker and observed the resulting performance metrics (PCC). Specifically, when the model was trained with GSC1 and GSC2 patient data was the evaluation dataset (GSC1→GSC2), the most striking change occurred when we perturbed the H3K27Ac signals. Here we saw a decline in performance of 0.583851 (70.667%) in PCC. We noted declines of 0.036298 (4.393%) for RNAPII, 0.004939 (0.597%) for ATAC-seq, and 0.00419 (0.507%) for CTCF perturbations (Fig 5A). This suggests that the predictive dependence of gene transcription was on all epigenetic markers (histone modifications, RNAPII binding, broad chromatin accessibility, and chromatin looping), with a greater weight on H3K27Ac signals followed by RNAPII, ATAC-seq, and then CTCF.

To determine if this finding is consistent across patients, we applied our perturbation methodology to the opposite prediction direction (GSC2→GSC1). The mean of all the experiments produced a decline in PCC, although to a lesser degree. The perturbation of the H3K27Ac signal of the GSC1 patient data while it was the evaluation dataset yielded a metric decrease of 0.14429 (17.824%). This was lower than the percentage decrease of the same feature as seen above, but it was a higher difference when compared to the other three features. The remaining results for this prediction direction showed declines of 0.008748 (1.08%) for RNAPII perturbation, 0.005978 (0.738%) for ATAC-seq, and finally 0.0021 (0.2594%) for CTCF perturbation (Fig 5B). This other direction of cross-patient in-silico perturbation analysis shows the same order of the predictive epigenetic modulator’s dependence on gene transcription is observed in the other patient. This suggests that, according to the data, gene transcription is highly connected with the H3K27Ac signals, followed by RNAPII, ATAC-seq, and CTCF. And this mechanism is observed across both patients. Interestingly, perturbing H3K27Ac led to different degrees of PCC reduction in GSC1→GSC2 and GSC2→GSC1 (0.456 and 0.250, respectively). This indicates that the degree of gene transcription dependence on H3K27Ac signals varies among GSCs. This could be potentially due to their heterogeneity, as H3K27Ac signals can vary among patients (29).

**Fig 5A & 5B.**
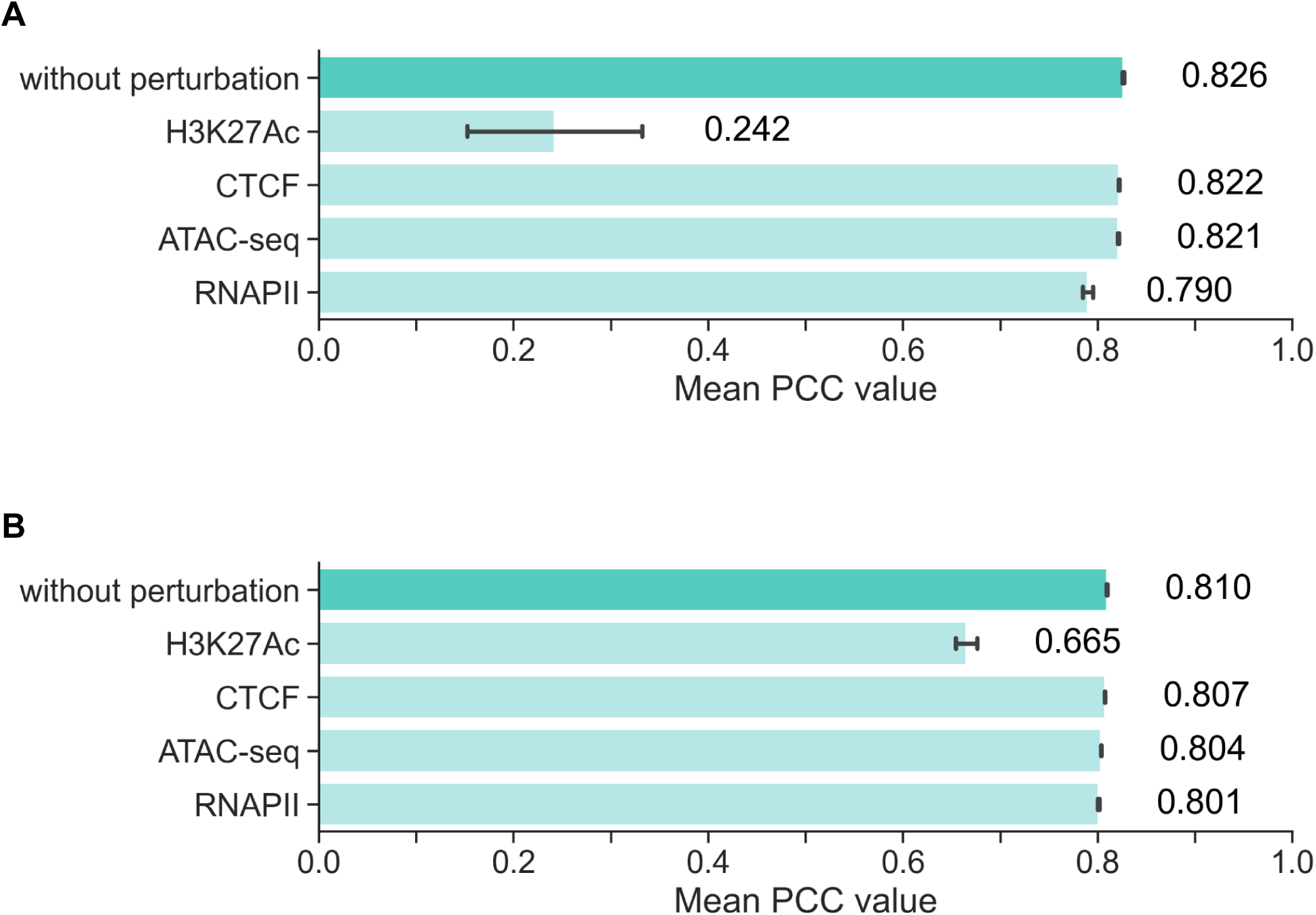
Epigenetic signal perturbation model performance comparison using XGBoost in both prediction directions. Each epigenetic signal was perturbed over 10 separate experiments in the two prediction directions. Shown are the mean PCC model results for 10 runs for each epigenetic marker (different random seeds) and error bars indicating the standard deviation. The hyperparameters used were identical to our other testing (see Table 1). The figures illustrate the order of model effect signal perturbation had from most to least: H3K27Ac, RNAPII, ATAC-seq, and CTCF. The effect was consistent in both prediction directions. A) GSC1→GSC2 prediction direction. B) GSC2→GSC1 prediction direction.

Comparing these PCC values to the correlation values between RNA-seq and each epigenetic marker, we can say that our cross-patient prediction analysis captured a trend that is not captured by simple correlation analysis (S4, Fig S3A & S3B). Additionally, this result on the importance of H3K27ac signals was furthered supported by the SCC results (S7, Fig S5A & S5B) and the primary model’s feature importance output (S8, Fig S6A & S6B). To address the generalibiilty of this model to unseen data, we altered the hyperparamters for the “reverse” direction, and observed a measurable increase in PCC/SCC, suggesting the generalibility of this model (S9).

### 4.3 Cross-patient analysis with neural crest and progenitor cell epigenetic data

H3K27Ac is known as an enhancer marker, and CTCF is also known as a mediator between enhancer and promoter. The enhancers are crucial gene expression modulators among GSCs as well as different cell types such as neural crest and progenitor cells. Thus, we hypothesized that H3K27Ac and CTCF would be critical markers to predict gene expression for GSCs as well as NCCs and neural progenitor cells. Furthermore, we hypothesized that if we test our GSC-trained model with neural crest or progenitor cells, we would observe similar PCC values. To test this hypothesis, we used publicly available data of H3K27Ac and CTCF-ChIP seq of neural crest and progenitor cell as test data and compared the PCCs. When we test the patient dataset on the trained model, we curated each dataset to include their H3K27Ac and CTCF values while ensuring that the ATAC and RNAPII values were 0 (the mean of the standardizations) across all genes of GSCs and then compared the PCCs. Our cross-cells analysis (Fig 6A and 6B) showed that testing the model with GSC showed the highest PCC (0.780-0.794) followed by one with NCC (0.649-0.672), one with neural progenitor cells (0.549-0.567). This indicates that computationally H3K27Ac/CTCF-related epigenetic landscape is most similar across GSCs, then followed by GSCs and NCCs, and GSCs and neural progenitor cells.

**Fig 6A & 6B.**
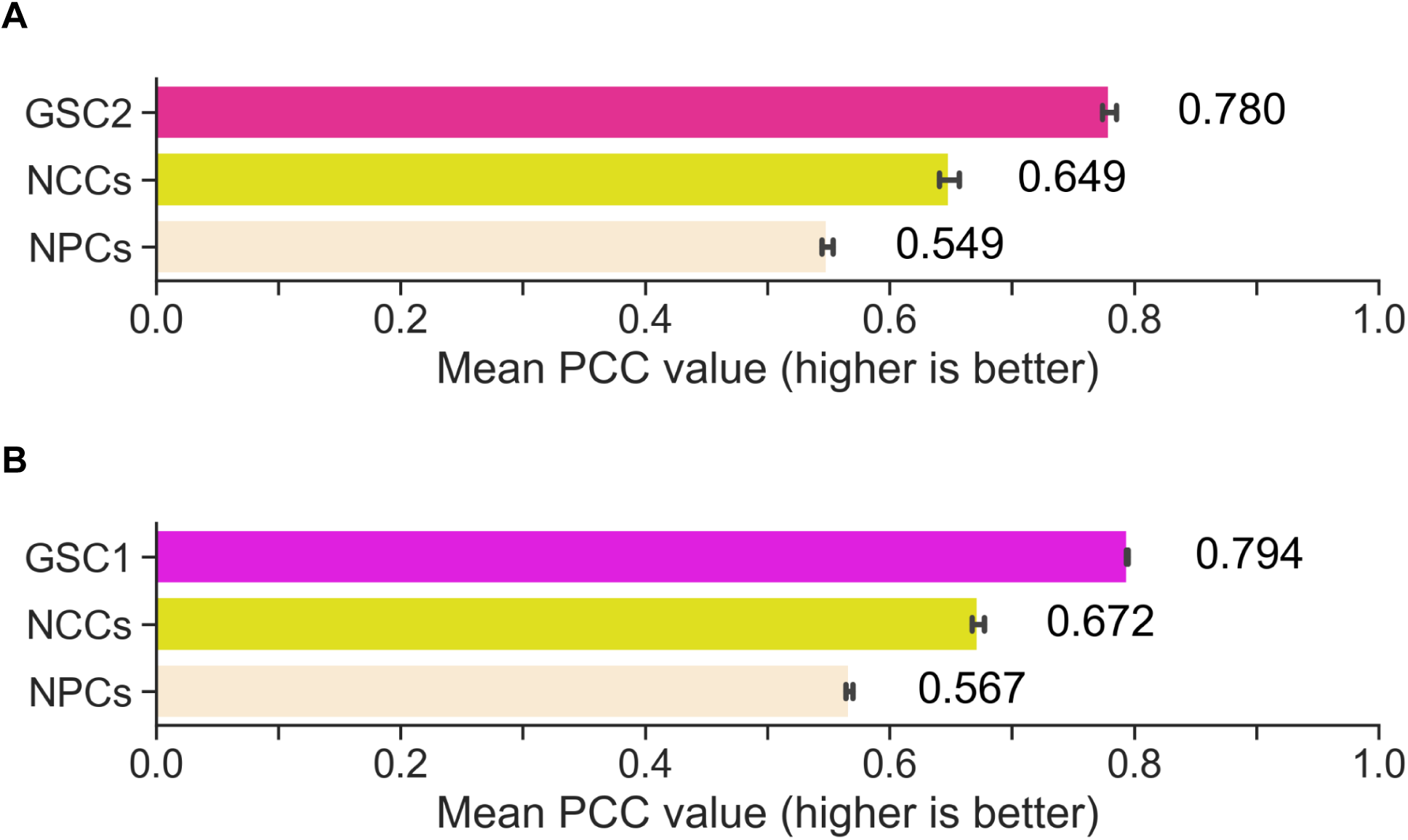
Our cross-patient analysis was extended to include neural crest and neural progenitor cell data compiled from features available on ENCODE. In each experiment, the test set included only the H3K27Ac and CTCF markers. A) The model was trained with GSC1. From the prediction performance it was inferred that the GSCs were most similar to each other. Interestingly, the plot indicates that the NCC epigenetic data is more similar to GSC1 than the neural progenitor cell data. B) When the model was trained with GSC2, we saw the same trend where the GSCs were most similar followed by the NCC data. Both plots’ values are the mean of 10 experiments and the standard deviation indicated by the error bars.

To further investigate this similarity in epigenetic landscape, we particularly looked into the genes with higher accuracy as these genes significantly contribute to training the model. Our XGBoost-trained model allowed us to rank genes based on prediction accuracy and to identify a group of genes that contributes to training the model. Thus, we ranked the genes based on performance, which is defined by means of squared error between observed and predicted values from 10 runs. Particularly, we focused on only expressed genes, because H3K27Ac and CTCF are known to positively correlate with gene expression, thus low-expressed genes would not be influenced by their H3K27Ac and CTCF signals around TSS (11,29). Looking at the rank of genes for all the GSC, neural crest and progenitor cells, we set 4500 as a threshold to separate between “accurately predicted” genes and the rest. Our PCC result indicates that GSCs are more similar to NCCs than to neural stem cells in terms of H3K27Ac-and CTCF-based epigenetic landscape. Given this data, we decided to focus on GSCs and NCCs.

To assess the epigenetic landscape of GSC1, GSC2, and NCCs, we looked into the intersection of “accurately predicted” genes between GSC1, GSC2, and NCCs and identified 544 genes in the intersection. Furthermore, we looked into the aggregated signals of these 544 genes for H3K27Ac and CTCF, respectively, and compared it to the aggregated signals of randomly selected genes (Fig 7A and 7B). For H3K27Ac, we observed a shift in the peak distribution for the “accurately predicted” genes of NCCs compared to the randomly selected genes, suggesting that H3K27Ac signals contribute to accurately predicting gene expression of NCCs. (Fig 7A), Given this result and H3K27Ac’s contribution on predicting GSC’s gene expression, we hypothesized that the intersection peaks for NCCs would show good concordance with the GSC peaks, and in fact it did. Meanwhile, considering the epigenetic disparities between GSCs and NCCs, as expected, we observed the small difference in standardized counts right before TSS (e.g. around bin 22-23) between GSCs and NCCs (Fig 7B). Regarding CTCF, we observed a comparable peak distribution between “accurately predicted” genes and randomly selected genes. This indicates that CTCF of “accurately predicted” genes don’t significantly contribute to predicting gene expression. Moreover, the contribution of each epigenetic marker, H3K27Ac and CTCF, of “accurately predicted” genes align with the observed difference in their predictive capabilities for gene expressions. Overall, this result suggests that H3K27Ac contributes to accurately predicting gene expression of NCCs as it does with GSCs. However, CTCF seems to be less impactful for prediction. Additionally, the resemblance of H3K27Ac peak distribution between GSC and NCC underscores epigenetic similarities at the enhancer landscape when it comes to predicting gene expression.

**Fig 7A & 7B.**
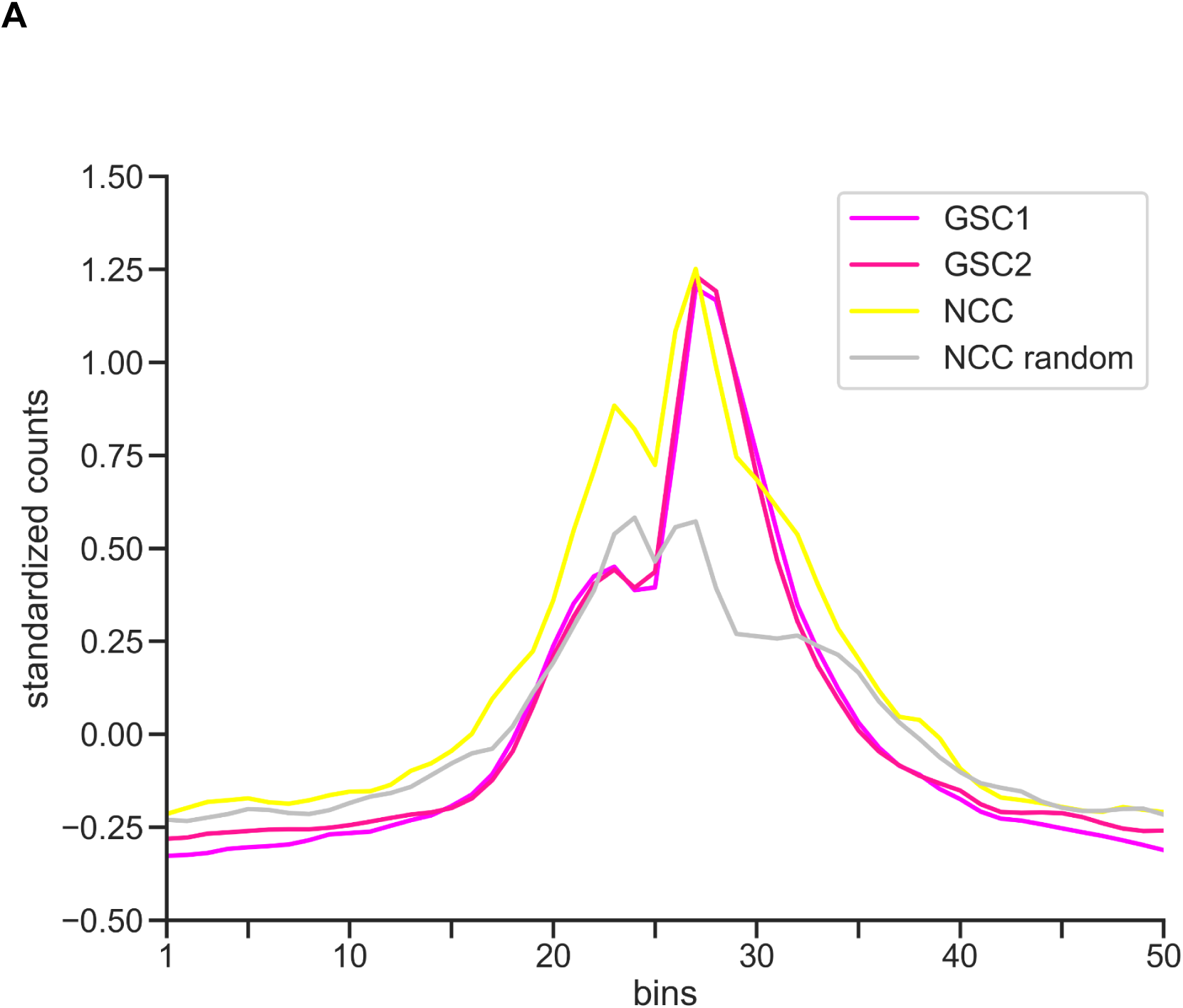
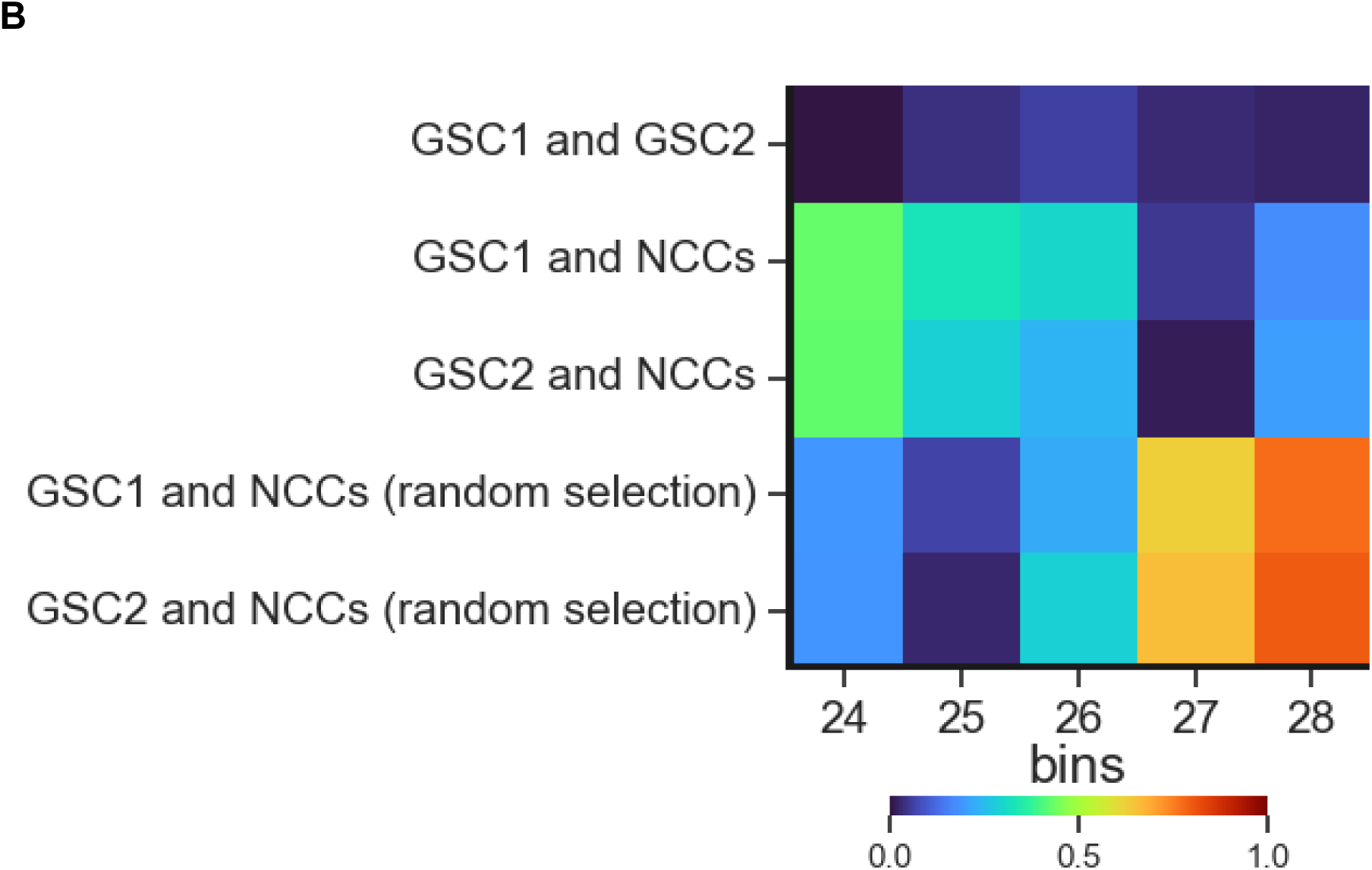
Epigenetic signal analysis. We performed an analysis of H3K27Ac signals (standardized counts per bin), to investigate similarity/dis-similarity between GSCs and NCCs. A) Each line represents the mean of the standardized count values for 544 common genes identified by the intersection of the model’s lowest error per gene, on the respective test set. The plot shows the bi-modal nature of the H3K27Ac signal with corresponding peaks at the bin 27 within the TSS region. B) The heatmap visualizes the Euclidean distance between pairs of dataset mean values in the highlighted region. We identified similarity when comparing GSC1 and GSC2 indicated by the relatively small distance between their mean values. The same observation of similarity was made at bin 27 where the Euclidean distance is also relatively small between the GSCs and the NCC datasets. The observation is contrasted by the relatively larger difference in the signals at bin 27 and 28 between the GSCs and a group of corresponding 544 genes within the NCCs that were randomly chosen from outside of the lowest error rate gene population.

## 5. Discussion

Origin and maintenance of GSC plasticity are regulated by endogenous cell processes affecting DNA, chromatin, and RNA, as well as by factors of the microenvironment that help propagate cancer stem cell phenotypes. Epigenetic pressure contributes to the ability of GSCs to remain plastic and allows GSCs and differentiated progenies to adopt a population equilibrium that facilitates tumor persistence (30–32).

To understand non-linearity of epigenetic mechanisms that drive gene expression, machine-learning has been recently employed as demonstrated in AttentiveChrome and GraphReg (16). Here, we apply machine learning to a combination of epigenetic bulk NGS data to discover epigenetic marks that can predict gene expression across patients with glioblastoma. To explore the developmental origin of cellular plasticity in GSCs, we successfully developed machine-learning models that utilize cross-patient prediction to investigate the epigenetic regulation of gene expression in heterogeneous GSCs, NCCs, and NPCs.

Our model comparison shows XGBR architecture to be the best performing model. XGBR is known as a strong architecture for processing tabular data and our datasets consisted of various types of epigenetic modulators arranged as a series of tables (33). Our cross-patient perturbation analysis using the XGBR model indicates that H3K27Ac signal contributes more to prediction of gene expression compared to RNAPII, broad chromatin accessibility, and chromatin looping across patients. Our perturbation analysis shows that the sum of the drop in PCC by each epigenetic modulator is less than PCC without perturbation. This suggests that other potential contributors to gene expression prediction may exist, such as other histone modification marks (e.g., H3K9me3, H3K27me3, etc.) and modulators of distal chromatin looping (e.g., Cohesin).

Recently, application of a neural network projection on the developmental trajectory of normal brain cells uncovered that glioblastoma cells share features of common lineage with perivascular neural crest and radial glial cells (34). Here we show that patients derived GSCs exhibit common patterns of H3K27Ac marks with NCCs. Remarkably, H3K27Ac marks of human NCCs can predict gene expression of GSCs from different patients with glioblastoma. Since these are bulk NGS data, they suggest a conserved enhancer landscape between certain subpopulations of GSCs and NCCs. In the future, it will be important to define the specific subpopulation of GSCs that shares gene regulatory networks with NCCs to determine conserved epigenetic traits of cellular plasticity between glioblastoma and the developing nervous system.

## 6. Conclusion

We built a cross-patient prediction analysis framework that can be used to provide insights on the contribution of multi-epigenetics markers on predicting gene expression of cancer stem cells and other cells with stem-cell phenotypic potential that involves cell plasticity. By applying this framework to GSCs, we identified that H3K27Ac signals contribute to predicting gene expressions most, followed by RNAPII, ATAC-seq, and CTCF, and this contribution is preserved across patients. Furthermore, we applied this to neural progenitor cells and crest cells and found that H3K27Ac/CTCF-related epigenetic landscape was similar across GSCs, neural progenitor cells, and neural crest cells. Overall, we presented a cross patient gene expression prediction framework that can be used to formulate deep insights into epigenetic-driven gene expression mechanisms and the epigenetic landscape of cellular plasticity across cancer stem cells and multiple cell types.

## 7. Abbreviations

GSC1: Glioblastoma Stem Cells 1. Nomenclature to indicate patient 1.
GSC2: Glioblastoma Stem Cells 2. Nomenclature to indicate patient 2.
GSC1→GSC2: This denotes the experimental setup whereby the model is trained on data derived from patient GSC1 (patient 1) while the test set is composed of data from patient GSC2 (patient 2).
GSC2→GSC1: Indicates that the training dataset is derived from patient GSC2 and the prediction direction is toward the test set of GSC1.
NCC: Neural Crest Cell
NPC: Neural Progenitor Cell
PCC: Pearson Correlation Coefficient metric.
SCC: Spearman Correlation Coefficient metric.
SD: Standard deviation.

## 8. Code availability

The study’s code is located at https://github.com/rsinghlab/ML_epigenetic_features_glioblastoma.

## Supporting information

### S1. Epigenetic marker and RNA-sequencing information

#### GSC datasets

##### Primary glioblastoma stem cell isolation and culture

Primary human glioblastoma stem cells (GSCs) were isolated from Glioblastoma Multiforme (GBM) tumors using an established selection media-based protocol. GSCs were cultured in complete culture media: Neurobasal-A (Fisher Scientific, 10888022), B27 minus Vitamin-A (Fisher Scientific, 12587010), Glutamax (Fisher Scientific, 35050-061), Heparin (StemCell Tech, 07980), human HB-EGF, 100 ug (Peprotech, 100-47), human FGF,100 ug (Peprotech, 100-18B), and Antibiotic: Antimycotic (Anti-Anti) (Gemini Bio-products, 400-101).

##### RNA-sequencing data

GSCs were lysed using the Trizol reagent (Invitrogen). RNA was isolated from this lysate using the RNeasy mini kit (Qiagen). Next-generation sequencing was performed on these RNAs.

##### RNA-pol II and H3K27Ac ChIP-sequencing data

For each biological replicate, 5 million cells were pelleted and sent to Active Motif for chromatin immunoprecipitation, library preparation, and bioinformatic analysis. 75-nucleotide sequence reads were obtained using the Illumina NextSeq 500 system, resulting in over 30 million reads for each library, and aligned to the genome using the Burrows-Wheeler Alignment (BWA) algorithm. The 3’ end of aligned reads (tags) were extended in silico to 150-250 bp and the density of these extended fragments was quantified for the entire genome, which was divided into 32-nucleotide bins. The detection of peaks, genomic regions with significant local enrichment for tags, was performed using the MACS and SICER algorithms. Tag number across multiple libraries was normalized by being reduced to the number of tags in the smallest library through random sampling, in order to preserve site-specific and global differences between libraries.

##### CTCF ChIP-seq

For each biological replicate, 3 million cells were crosslinked and lysed using the truCHIP Chromatin Shearing Kit (Covaris). Isolated chromatin was sheared at 105 peak incident power, 2% duty factor, and 200 cycles per burst for 100 seconds in an S220 Focused-ultrasonicator (Covaris) to a length of 200-1200 bp. Shearing efficiency was validated using a Fragment Analyzer (Agilent). Immunoprecipitation for CTCF and ChIP-seq library preparation was performed using the ChIP-IT High Sensitivity Kit (Active Motif) and Next Gen DNA Library Kit (Active Motif), following the manufacturer’s instructions.

##### ATAC-seq

For each biological replicate, 50,000 cells were used to prepare ATAC-seq libraries in accordance with the protocol used in Ackermann et al. for alpha and beta cells, which was adapted with some modifications from the original protocol from Buenrostro et al. and the Omni-ATAC protocol. Samples with low viability (85%) were treated with DNase (Worthington) at a concentration of 200 U/mL for 30 minutes at 37° C. Double-sided bead purification to remove primer dimers and large 1,000 bp fragments was carried out using Agencourt AMPure XP beads (Beckman Coulter). Quality control was performed before sequencing using a Fragment Analyzer (Agilent) and the KAPA Library Quantification Kit (Roche). Libraries were analyzed by GENEWIZ using the Illumina HiSeq 2500 system to acquire 150 bp paired-end sequence reads. 200 million genomic reads per sample were obtained to detect open vs. closed chromatin regions.

#### Neural Progenitor Cell Dataset

Our progenitor cell datasets were composed from marker bam alignment files available from ENCODE (https://www.encodeproject.org) (20–26,35). They were all derived from the biosample ENCBS018TPT. Since RNAPII ChIP-sequencing was unavailable, we used the signal value of 0.000000 (the mean of the standardization) across all gene bins.

##### RNA-sequencing data

This RNA-seq data was from the lab of Thomas Gigeras, CSHL under experiment ENCSR244ISQ, We chose isogenic replicate 1 which was from accession ENCFF056WDN (https://www.encodeproject.org/files/ENCFF056WDN).

##### H3K27Ac ChIP-sequencing data

This data was listed under accession ENCFF521XJN (https://www.encodeproject.org/files/ENCFF521XJN) as isogenic replicate 2. It was from the lab of Bradley Bernstein, Broad from experiment ENCSR449AXO.

##### CTCF ChIP-sequencing data

For this feature we chose isogenic replicate 2 in accession ENCFF400MZX (https://www.encodeproject.org/files/ENCFF400MZX). lENCODE lists the experiment as ENCSR125NB from the Lab of Bradley Bernstein, Broad.

##### DNase-sequencing data

For the progenitor cell dataset we used DNase-seq data as an analog for ATAC-seq. This particular data was from the lab of John Stamatoyannopoulos, UW. It was listed under ascension ENCFF123YLB (https://www.encodeproject.org/files/ENCFF123YLB) as isogenic replicate 1 and was from the same biosample (ENCBS018TPT) as the other data for this dataset.

#### Neural Crest Cell Dataset

The neural crest cell dataset was composed of H3K27Ac, CTCF and RNA-seq marker data from ENCODE (20–26,35). They were all derived from the ENCBS417JEU biosample. The features that were unavailable, RNAPII ChIP-sequencing and ATAC-seq, were replaced with the value 0.000000 across the bins of each gene.

##### RNA-sequencing data

This dataset’s RNA-seq values were from experiment ENCSR761SHI, attributed to the lab of Barbara Wold, Caltech. The isogenic replicate used was replicate 2 under the accession of ENCFF503KKJ (https://www.encodeproject.org/files/ENCFF503KKJ).

##### H3K27Ac ChIP-sequencing data

Our study used the data from accession ENCFF655GGB (https://www.encodeproject.org/files/ENCFF655GGB) which was isogenic replicate 1. The data was collected under experiment ENCSR685HSP from the lab of Bradley Bernstein, Broad.

##### CTCF ChIP-sequencing data

This CTCF data was also from the lab of Bradley Bernstein, Broad. In this case it was from experiment ENCSR218MVT which included isogenic replicate 2. It can be found under accession ENCFF583OOM on ENCODE (https://www.encodeproject.org/files/ENCFF583OOM).

### S2. Flowcharts of data preparation and preprocessing process

Our patient dataset preparation process involves a number of utilities which convert the feature reads housed in BAM files into counts for each of 50 bins for all 20,015 genes. As illustrated in Figure S1, each of the four epigenetic signal data collections follow one preparation path while the expression data follows another. Ultimately, the two preparation paths converge to produce 2-dimensional, 1,000,750 x 5 files for model script input.

Figure S2 depicts our data preprocessing just prior to model input within the script. Again, we saw that the epigenetic feature and target variable follow different paths for their respective preprocessing. The feature data leading to the dataset splitting process before standardization. Meanwhile, the target variable undergoes a log(2) transformation before the appropriate dataset splitting. The processes were coordinated to ensure that each gene’s information remains synchronized throughout.

Given that all of our models were configured for cross-patient prediction, the data preprocessing occurs in sequence for each patient datafile as the model script is run.

**Fig S1.**
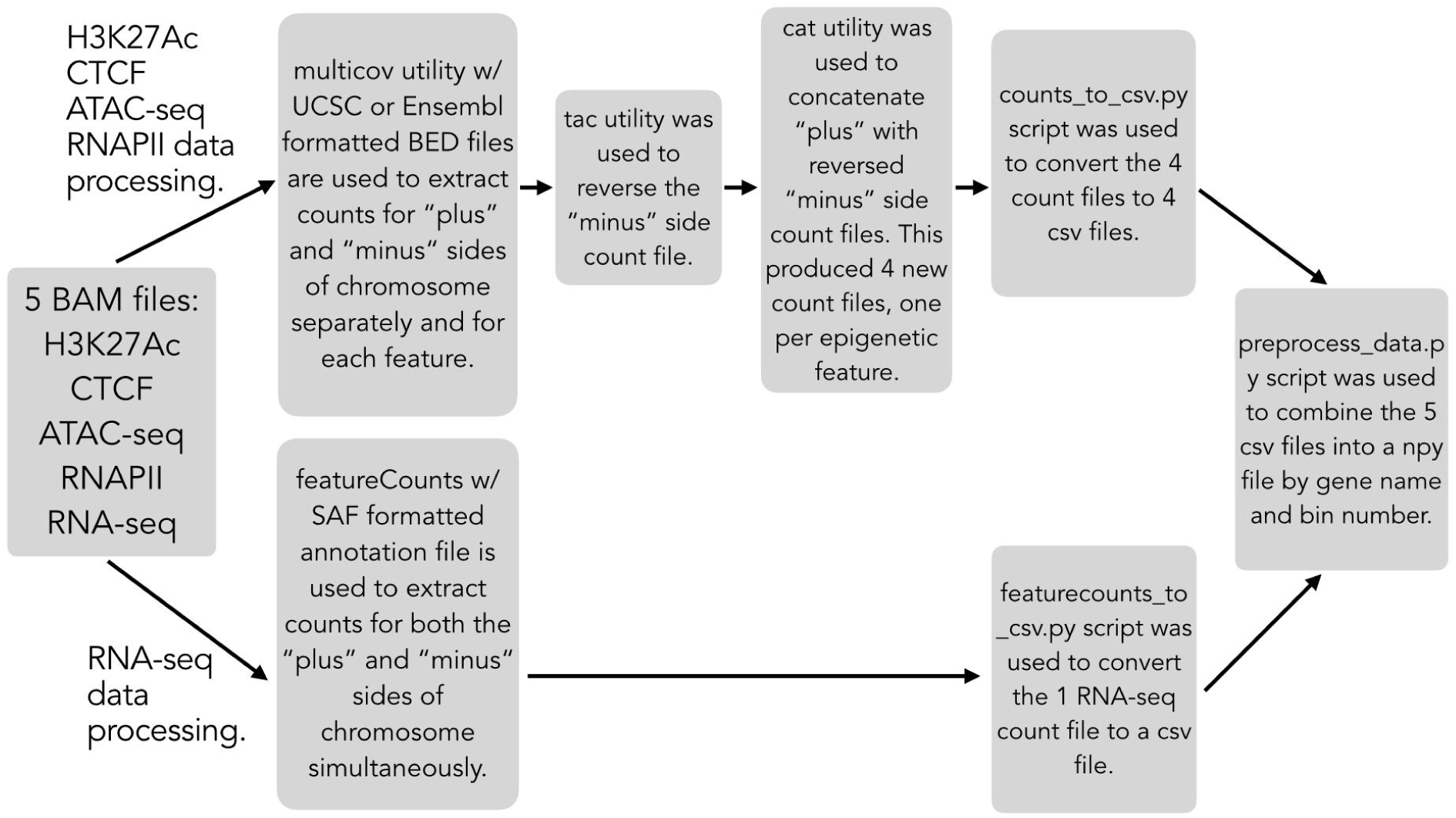
Illustration of dataset preparation process. The process that extracted the observed epigenetic marker values from BAM format files made use of a series of utilities which combined the separate data into one file. The process aligns the 50 bin values of the gene’s epigenetic features with a single extracted RNA-seq value for the gene to maintain consistent formatting.

**Fig S2.**
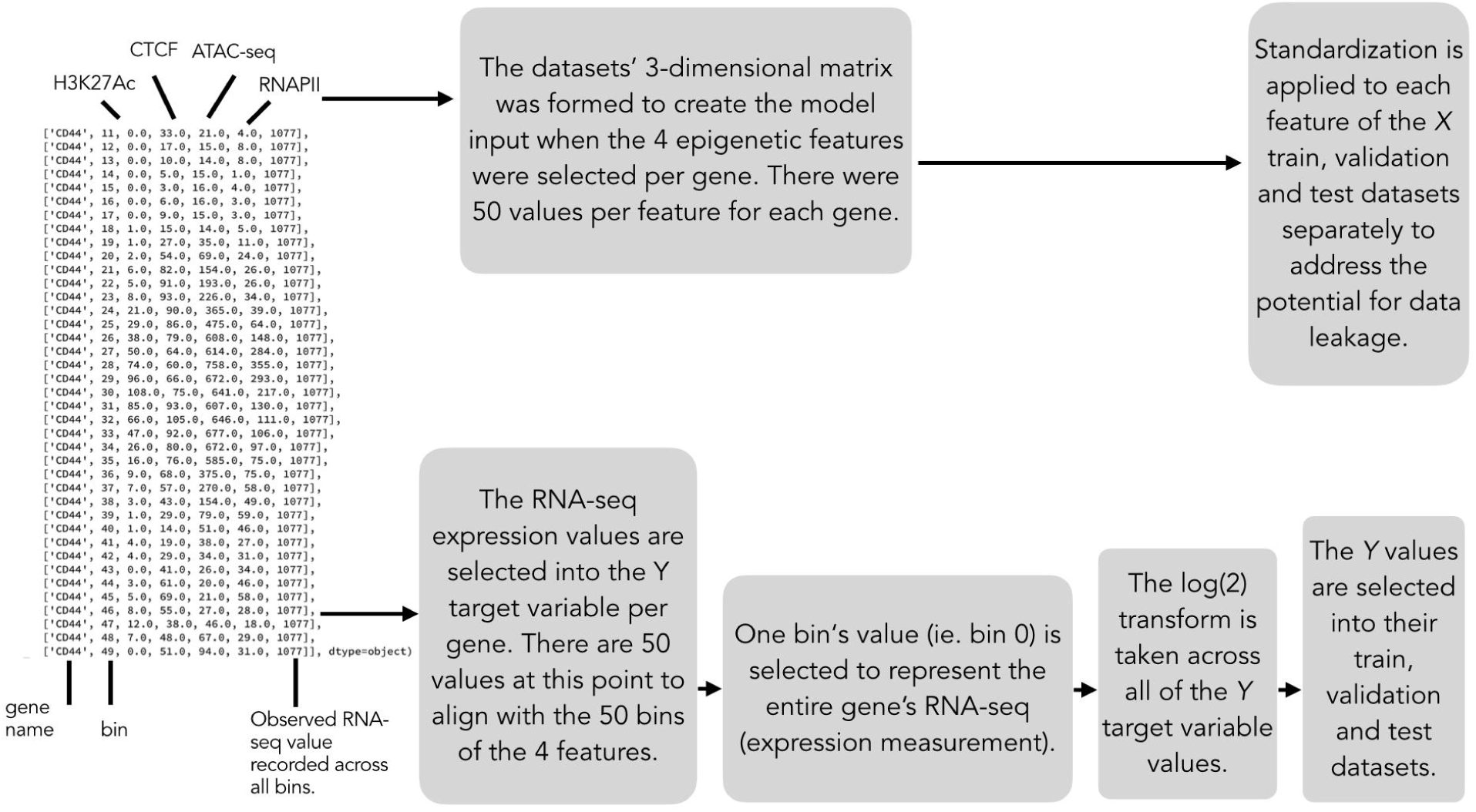
Illustration of data preprocessing process. The epigenetic feature values per gene were extracted, formed into matrices, and finally standardized for model input. Each gene’s observed RNA-seq value is extracted and transformed creating the target variable for the model.

### S3. Supporting predictive models information

#### XGBoost Regression (XGBR)

An overview of the parameters we tuned starts with η (learning rate), which referred to the step size shrinkage of the feature weights at each iteration. "Max depth" referred to the maximum depth for each tree while "n estimators" determined the total number of trees in the model. γ (min split loss) is the minimum reduction in loss required to split on a leaf node. A parameter that the algorithm used to reduce overfitting is "subsample" or the ratio of random samples taken from the training data before growing a new tree. The algorithm used "colsample bytree" which was a ratio to determine the number of features to be randomly selected for each tree. Meanwhile, "min child weight" referred to the minimum number of samples (the weight) required to form a new tree node(36,37).

**Table S1.** XGBoost Regression model hyperparameters. Outlined in the table are the chosen hyperparameter values from their respective search spaces over our tuning process.

| hyperparameter | model value | search set |
| --- | --- | --- |
| $\eta$ | 0.03 | 0.01, 0.02, 0.03, 0.04, 0.05, 0.07, 0.10, 0.12, 0.15, 0.17, 0.20, 0.25, 0.30, 0.40, 0.50 |
| max depth | 6 | 2, 3, 4, 5, 6, 7, 8, 9, 10, 12, 15, 17, 20, 25 |
| number of estimators | 150 | 25, 50, 75, 100, 110, 125, 130, 150 |
| $\gamma$ | 0.3 | 0.0, 0.1, 0.3, 0.5, 0.7 |
| subsample | 1.0 | 0.2, 0.4, 0.6, 0.8, 1.0 |
| column sample by tree | 0.2 | 0.2, 0.4, 0.6, 0.8, 1.0 |
| minimum child weight | 1 | 1, 3, 5, 7, 9 |

#### Multi-layered Perceptron (MLP)

Our Multi-layered Perceptron model was found to be one of our most performant configurations for testing and prediction in our cross-patient experiments. The 3-dimensional dataset structure was kept intact prior to model input. Each gene’s epigenetic features were then flattened at the next layer. Since batching was used, the dataset matrix went from *b* x 50 x 4 to *b* x 200 where *b* represented the batch size. The model calculations then progressed through a collection of three hidden (dense) layers alternating with two dropout layers (for regularization). The model’s 4th dense layer is configured with a single output (with linear activation) for RNA-seq value prediction for this regression task.

This model’s hyperparameters tuning followed the same methodology as our other models. Each of the three dense layer sizes were tuned independently of each other. Our tuning process resulted in an unconventional configuration where the number of nodes in a layer downstream from previous ones is higher than the others. We then experimented with fine tuning the layer sizes relative to each other and found that the existing setup performed well. Additionally, we opted to configure the dropout rate as part of our grid search for tuning.

This model’s results were consistently near the highest we observed with our data. Going further, it was the best performing of our deep learning based architectures. Because of that we used the same experimental setup in a version of our perturbation analysis.

**Table S2.** Multi-layered Perceptron hyperparameters.

| hyperparameter | model value | search set |
| --- | --- | --- |
| learning rate | 0.0001 | 0.01, 0.05, 0.001, 0.005, 0.0001, 0.0005 |
| $b$ | 512 | 512, 1024 |
| hidden layer 1 size | 100 | 20, 50, 100, 200 |
| hidden layer 2 size | 100 | 20, 50, 100, 200 |
| hidden layer 3 size | 200 | 20, 50, 100, 200 |
| dropout rate | 0.4 | 0.3, 0.4, 0.5 |
| epochs | 150 | Established by examining PCC metric training and validation plot curves. |

#### Branched Multi-layered Perceptron (Branched MLP)

Multi-modal computation was a significant portion of development phases and for a series of experiments we added an approach where we gave a model both the epigenetic features we discussed in the main body of this work along with the genetic sequence for each gene. The HG38 Human Reference Genome was used to create the sequence branch input. We acknowledged from the onset that the human reference genome may not be an informative source to partner with our glioblastoma epigenetic measurements since they are derived from different cell types. We proceeded with the experiments to determine if we would learn more about the model behavior and ability to predict gene expression.

As with our process for the epigenetic data, we developed an approach to the data that can be seen as having 2 parts; one internal and one external to the model script. The preparation process of the gene sequence data, outside of the script, incorporated the use of the same BED location files we used in our epigenetic data preparation process. The goal was to ensure that the location for sequence for each gene exactly matched the same its epigenetic markers. We collected 100 base pairs, ranging from above and below the gene’s TSS site, for each of 50 bins. Our next task was to convert the letter-based sequence to a numerical one. We used the following replacement strategy for this:

A = 1, T = 2, C = 3, G = 4, N = 5

The numpy files that resulted from these steps became additional input into the script where the second portion of the preprocessing took place. Here, the sequence data was composed 3-dimensionally where the gene sequence’s index position in this dataset was an exact match to its corresponding epigenetic measurement position. At this point we sought to consider the computational complexity of this branch of the model while maintaining the analysis of the data around the TSS site. The number of bins was reduced to 21 whereby, with TSS located at bin 25 the sequence input ranged from bins 15 to 35. Therefore, this led to an intermediate matrix shape of 20,0015 x 21 x 100. Finally, just prior to this data being input into the training/validation and evaluation model fits, our script one-hot encodes the numerical representations in the following way:

1 = [1, 0, 0, 0, 0], 2 = [0, 1, 0, 0, 0], 3 = [0, 0, 1, 0, 0], 4 = [0, 0, 0, 1, 0], 5 = [0, 0, 0, 0, 1]

The patient epigenetic measurement process was unchanged for this architecture so this model then received batched input for its respective “epigenetic” and “sequence” branches. Both branches have 3 dense layers with dropout layers in place for independent regularization. The output of those two branches was concatenated for input into a set of 4 dense layers. The first 3 layer sizes are tuned along with those in the aforementioned branches. The final dense layer is a single output for gene expression (RNA-seq) prediction.

In common with our other deep learning based architectures, this model used our measured RNA-seq values as the target variable, MSE for loss calculations, and the Adam optimizer. Additionally, hyperparameter tuning followed the same process as our other models.

Our Branched Multi-layered Perceptron performance was generally improved over the Multiple Linear Regression and Support Vector Regression. In turn, it was less performant than all other models. We also noted that its metric standard deviation was increased over all other models which may indicate that its cross-patient predictions are less stable than our other models.

Although we found other predictive algorithms to be more informative in this work, multi-modal approaches like this one are interesting for future investigation.

**Table S3.** Branched Multi-layered Perceptron hyperparameters.

| hyperparameter | model value | search set |
| --- | --- | --- |
| learning rate | 0.01 | 0.01, 0.05, 0.001, 0.005, 0.0001, 0.0005 |
| *b* (batch size) | 512 | 512, 1024 |
| epigenetic branch hidden layer size 1 | 200 | 20, 50, 100, 200 |
| epigenetic branch hidden layer size 2 | 200 | 20, 50, 100, 200 |
| epigenetic branch hidden layer size 3 | 200 | 20, 50, 100, 200 |
| epigenetic branch dropout rate | 0.3 | 0.3, 0.4, 0.5 |
| sequence branch hidden layer size 1 | 200 | 20, 50, 100, 200 |
| sequence branch hidden layer size 2 | 200 | 20, 50, 100, 200 |
| sequence branch hidden layer size 3 | 100 | 20, 50, 100, 200 |
| sequence branch dropout rate | 0.3 | 0.3, 0.4, 0.5 |
| combined branch hidden layer size 1 | 50 | 20, 50, 100, 200 |
| combined branch hidden layer size 2 | 200 | 20, 50, 100, 200 |
| combined branch hidden layer size 3 | 20 | 20, 50, 100, 200 |
| combined branch dropout rate | 0.3 | 0.3, 0.4, 0.5 |
| epochs | 150 | Established by examining PCC metric training and validation plot curves. |

#### Convolutional Neural Network (CNN)

We applied various configurations of Convolutional Neural Network (CNN) model architectures to our cross-patient prediction task as well. To accomplish this, we made use of Tensorflow/Keras model Application Programming Interface (API). As with our MLP model we used MSE for loss calculations along with Adam for optimization. The CNNs and MLPs also have the use of Dropout layers for regularization, in a number of places in their architectures. Our team’s experiments included 1, 2, and 3 layer CNN architectures; all built using Conv1D layers. In each case, pooling was implemented (MaxPooling1D layers). All of them fed their convolutional module output to a series of dense layers which lead to final prediction output. The layers used ReLU activation although we ran various experiments using LeakyReLU. Of note is that the 1 layer results were reported in the main body of this work by virtue of the fact that it received the most extensive tuning, training and testing within this group of models.

The nature of CNN layers and batching when fitting the model, allowed us to keep the datasets in their 3-dimensional structure for input after preprocessing. That made flattening only necessary just prior to the series of dense layers.

**Table S4.**
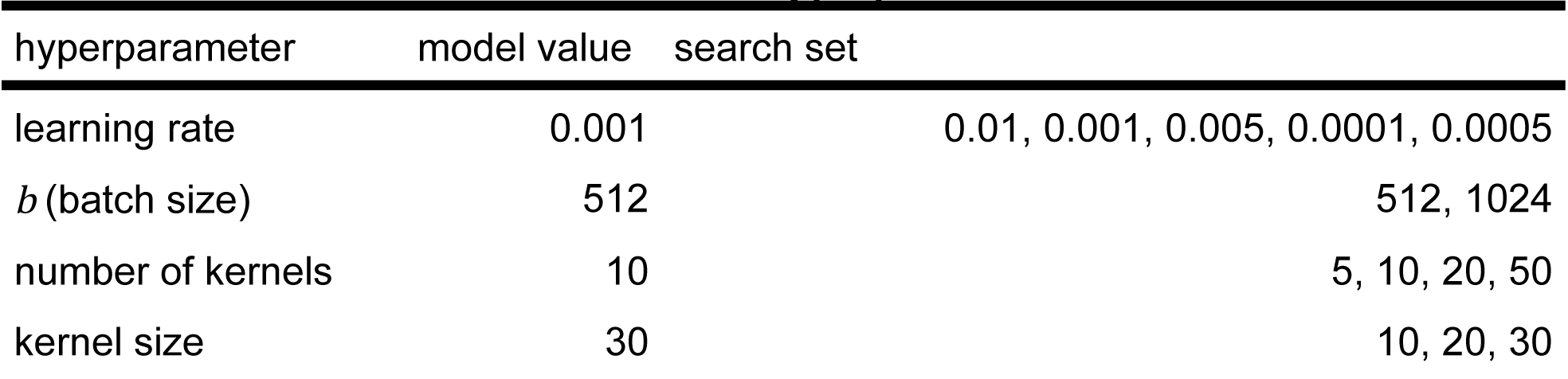
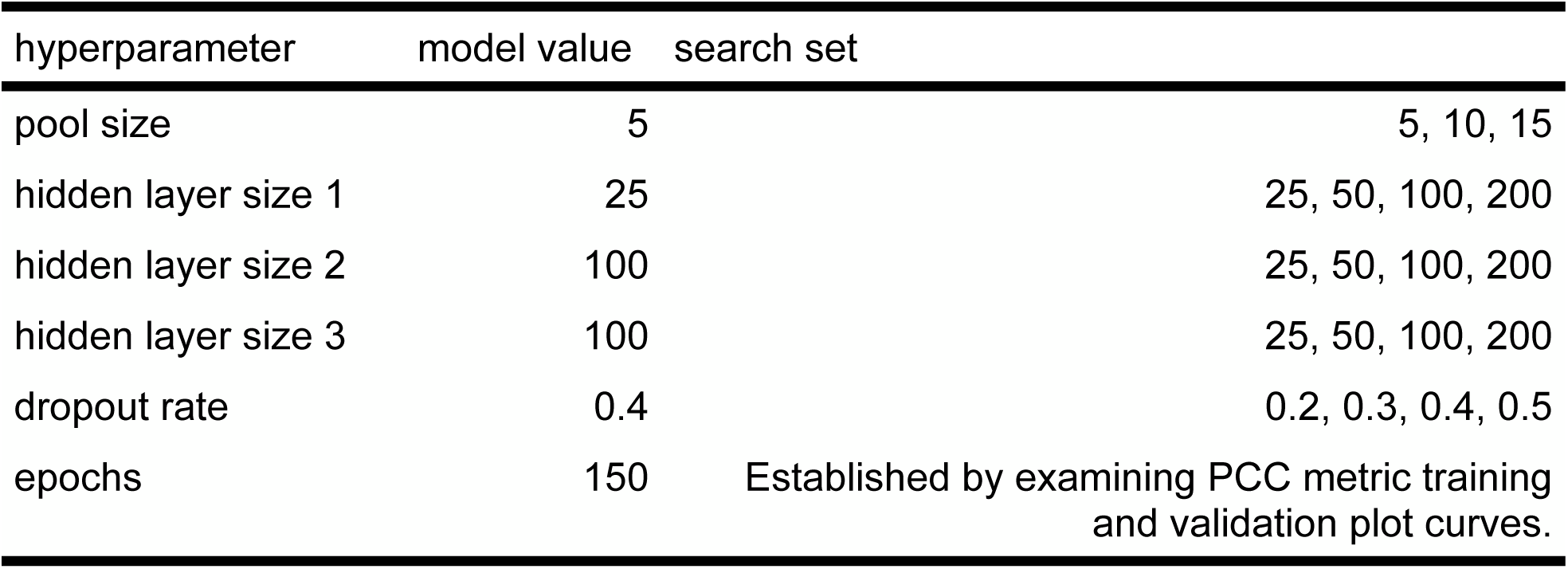
Convolutional Neural Network hyperparameters.

#### Recurrent Neural Network (RNN)

The sequential nature of the epigenetic features made Recurrent Neural Networks (RNNs) a suitable model choice for their capability to capture spatial relationships within the sequence. We conducted experiments with various RNNs and their extensions, and determined that AttentiveChrome, based on Gated Recurrent Units (GRUs) instead of Long Short-Term Memory (LSTM) (13), achieved the best performance on the dataset.

The AttentiveChrome architecture incorporates two levels of RNN and an attention mechanism. The first level consists of four GRUs, each processing information about one epigenetic feature. The input to these GRUs is treated as a time series, with each element in the sequence (i.e., each bin value for each epigenetic feature of a single gene) representing a time step (*x_t_*), with *t* ranging from 1 to *T* (*T*=50, as there are 50 bins). GRUs, as all other RNNs, maintain an internal state called the hidden state (*h*), which captures the historical information of the input sequence up to the current step. This hidden state acts as a form of memory, enabling GRUs to retain information about the observed data in the sequence and capture temporal (in terms of epigenetic features, spatial) dependencies. At each time step, the GRUs update their hidden state (*h*_t_) by considering the current input (*x*_t_) and the hidden state from the previous step (*h*_t−1_). GRUs employ two activation functions, functioning as an update gate and a reset gate, respectively, to selectively retain and erase information in the hidden state, mitigating the issue of vanishing gradients.

To enhance the GRUs’ ability to capture long-term dependencies and focus on relevant segments of the input sequence, a soft attention layer was applied to the hidden states of all time steps (*h*_1_*h*_2_…*h*_T_) from each of the four GRUs. The attention mechanism assigned weights to the hidden states, reflecting their relative importance, and thus added on the complexity and interpretability. The results are then concatenated and subsequently passed through a new GRU layer (with 4 time steps corresponding to the 4 epigenetic features), an additional attention layer, and a dense layer to generate the final predicted gene expression value. TensorFlow Keras was employed for the implementation of this model.

**Table S5.** Recurrent Neural Network hyperparameters.

| hyperparameter | model value | search set |
| --- | --- | --- |
| learning rate | 0.001 | 0.01, 0.05, 0.001, 0.0001 |
| $b$ (batch size) | 512 | 512, 1024 |
| RNN output size | 50 | 5, 10, 20, 30, 50, 80 |
| dropout rate | 0.1 | 0.1, 0.3, 0.5 |
| epochs | 200 | 100, 200, 300, 500 |

#### Gradient Boosting Regression (GBR)

Our first regression tree model configured for our cross-patient prediction analysis used scikit-learn’s library. Its performance after tuning was comparable with our deep learning models and was an early indication of the potential of tree based algorithms for prediction with our epigenetic patient data.

In the main body of this work, we briefly discussed our thoughts of the computational efficiency of our models. The library lacks graphical processing unit (GPU) acceleration and although the run times we recorded were competitive with the other models, this led us to consider the application of similar algorithms with GPU acceleration available.

**Table S6.**
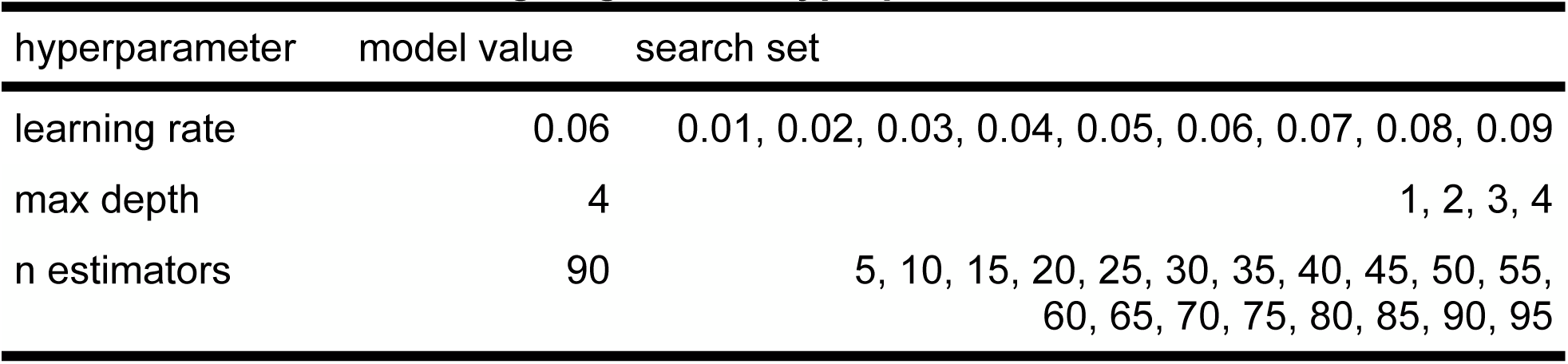
Gradient Boosting Regression hyperparameters.

| hyperparameter | model value | search set |
| --- | --- | --- |
| learning rate | 0.06 | 0.01, 0.02, 0.03, 0.04, 0.05, 0.06, 0.07, 0.08, 0.09 |
| max depth | 4 | 1, 2, 3, 4 |
| n estimators | 90 | 5, 10, 15, 20, 25, 30, 35, 40, 45, 50, 55, 60, 65, 70, 75, 80, 85, 90, 95 |

#### Support Vector Regression (SVR)

Our approach to building a regression version of the Support Vector Machine algorithm found us utilizing the scikit-learn library’s implementation together with our use of multiple patient datasets for model training and testing. Again, our datasets were reshaped, for model input using the identical method to our other traditional machine learning algorithm setups.

Our tuning phase centered on C and ɛ hyperparameters. Out of our search sets, we found 0.01 to be the optimal valve for both.

Across all of our testing, the SVR outperformed linear regression but did not reach the levels of performance reached by other models.

**Table S7.** Support Vector Regression hyperparameters.

| hyperparameter | model value | search set |
| --- | --- | --- |
| C | 0.01 | 0.01, 0.02, 0.03, 0.04, 0.05, 0.06, 0.07, 0.08, 0.09 |
| $\epsilon$ | 0.01 | 0.001, 0.002, 0.003, 0.004, 0.005, 0.006, 0.007, 0.008, 0.009, 0.01, 0.011, 0.012, 0.013, 0.014, 0.015, 0.016, 0.017, 0.018, 0.019, 0.02, 0.021, 0.022, 0.023, 0.024, 0.025, 0.026, 0.027, 0.028, 0.029, 0.03, 0.031, 0.032, 0.033, 0.034, 0.035, 0.036, 0.037, 0.038, 0.039, 0.04, 0.041, 0.042, 0.043, 0.044, 0.045, 0.046, 0.047, 0.048, 0.049, 0.05, 0.051, 0.052, 0.053, 0.054, 0.055, 0.056, 0.057, 0.058, 0.059, 0.06, 0.061, 0.062, 0.063, 0.064, 0.065, 0.066, 0.067, 0.068, 0.069, 0.07, 0.071, 0.072, 0.073, 0.074, 0.075, 0.076, 0.077, 0.078, 0.079, 0.08, 0.081, 0.082, 0.083, 0.084, 0.085, 0.086, 0.087, 0.088, 0.089, 0.09, 0.091, 0.092, 0.093, 0.094, 0.095, 0.096, 0.097, 0.098, 0.099 |

#### Multiple Linear Regression (MLR)

Our linear regression model was implemented using the statsmodels Application Programing Interface (API) where we chose the Ordinary Least Squares (OLS) version of the algorithm. Given that this is a traditional machine learning algorithm, our 3-dimensional datasets were reshaped into 2 dimensions before model input.

We chose to employ elastic net regularization when the model was fit to the data. This fit method allowed us to tune both the alpha (penalty weight) and L1_wt (the portion of the penalty applied to the L1 term).

**Table S8.** Multiple Linear Regression hyperparameters.

| hyperparameter | model value | search set |
| --- | --- | --- |
| $\alpha$ | 5000 | 0, 1, 5, 10, 25, 50, 75, 100, 200, 400, 800, 1600, 2500, 3200, 5000 |
| L1_wt | 0.0 | 0.0, 0.1, 0.2, 0.3, 0.4, 0.5, 0.6, 0.7, 0.8, 0.9, 1.0 |

### S4. Correlation highlights the variation among the epigenetic features for each dataset

To see if the application of machine and deep learning is appropriate to model the relationship between these epigenetic sequencing data and RNA-seq, we performed correlation analysis between gene expression and each epigenetic modulator (ATAC-seq, H3K27Ac ChIP-seq, CTCF ChIP-seq, and RNAPII ChIP-seq). To accomplish this, our patient datasets were each prepared in a similar fashion to what occurs in our models with some notable differences. The RNA-seq values were not log(2) transformed and the epigenetic sequencing data were not standardized. We instead calculated the sum of the bin counts for each gene’s 4 features where each row represented a gene’s bin position.

Our results indicated a positive correlation between gene transcription and each epigenetic modulator, with PCC values of 0.366, 0.111, 0.208, and 0.333 (for H3K27Ac, CTCF, ATAC-seq, and RNAPII, respectively) in GSC1, and PCC values of 0.412, 0.162, 0.221, and 0.359 (for the same order of modulators) in GSC2 (Fig S3A & S3B). These findings provided a promising starting point for exploring the potential of ML/DL models in this domain, based on the non-linearity between each modulator and RNA-seq which more complex machine and deep learning models can investigate.

**Fig S3A & S3B.**
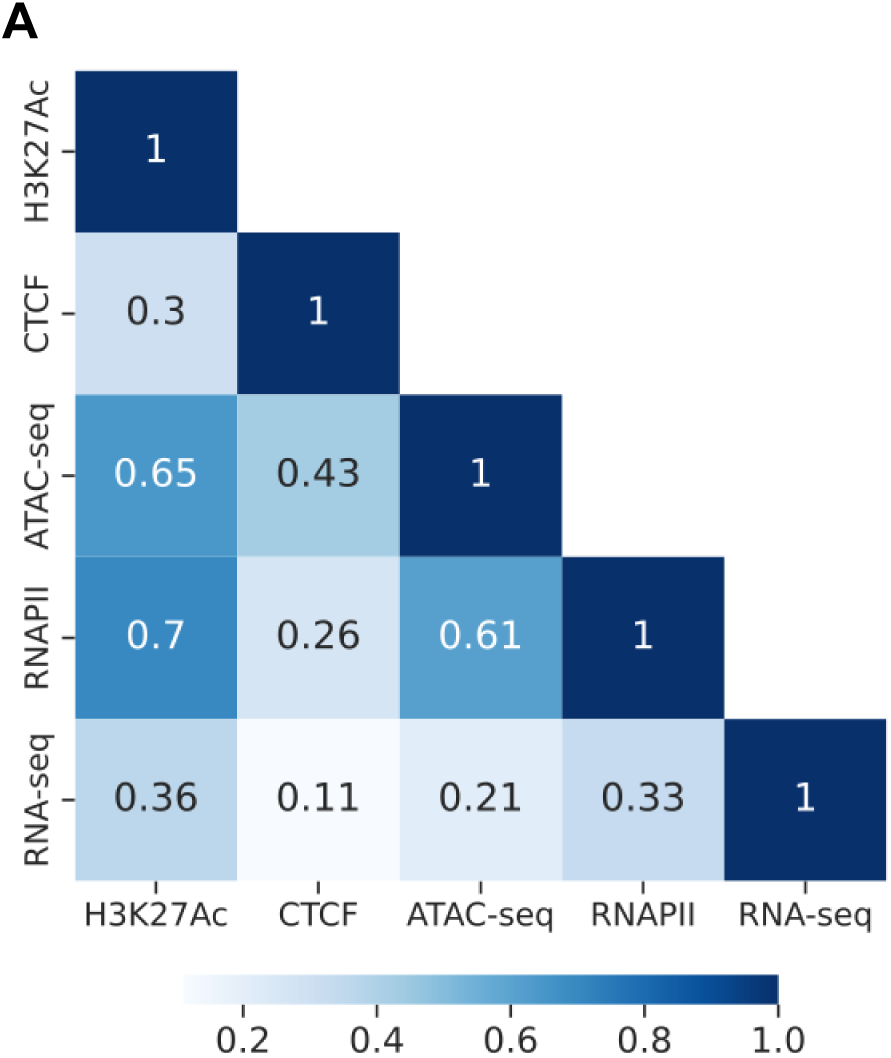
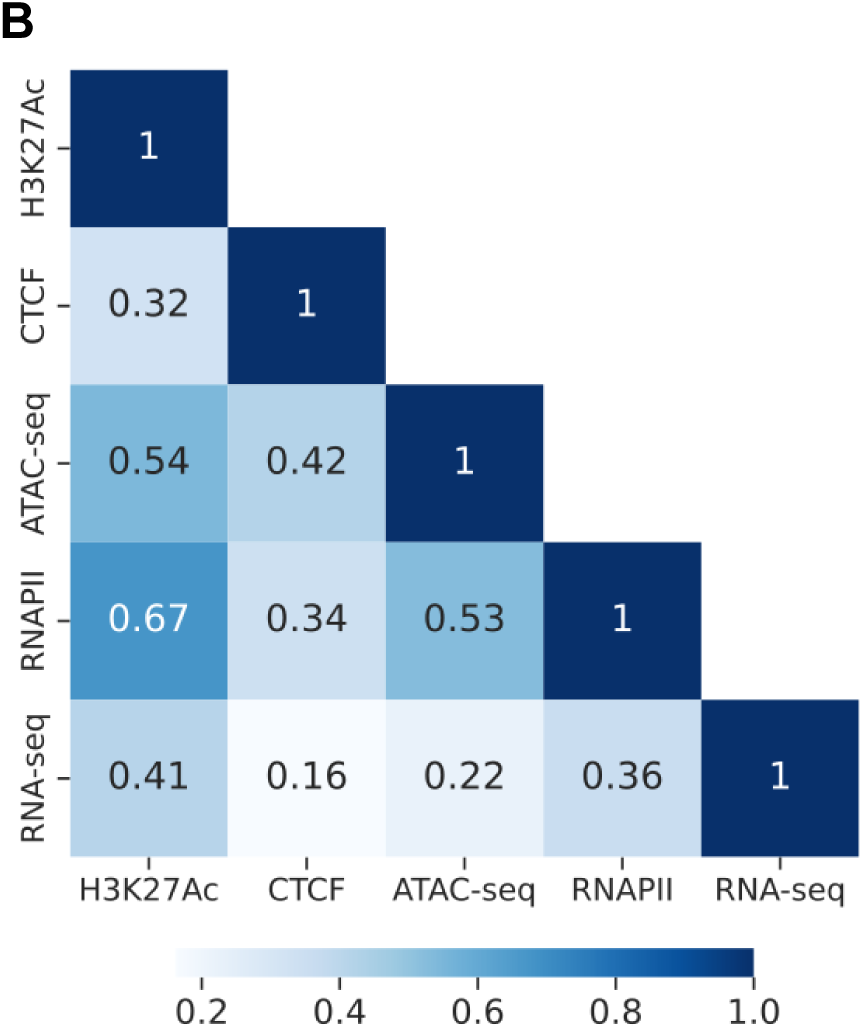
Correlation within GSC1 and GSC2 datasets. The two visualizations convey the calculated correlation values for our two patient datasets. GSC1 is shown in panel **A** with GSC2 in panel **B**. In both datasets the values indicate nonlinearity between the epigenetic modulators and gene expression.

### S5. Supporting model SCC results

All of our experiments produced Spearman Correlation Coefficient (SCC) results along with PCC. We designated it as our secondary evaluation metric and during our analysis, we noted that while many models’ values were comparatively lower than the corresponding PCC findings, the trends in how the models perform relative to each other remained the same. Our most performant model under PCC, XGBR, had higher results under SCC as well. It should be noted here that we sought hyperparameter combinations that optimized PCC and not SCC. With that in mind, it was interesting that the results we saw reinforce our PCC observations.

**Fig S4A & S4B.**
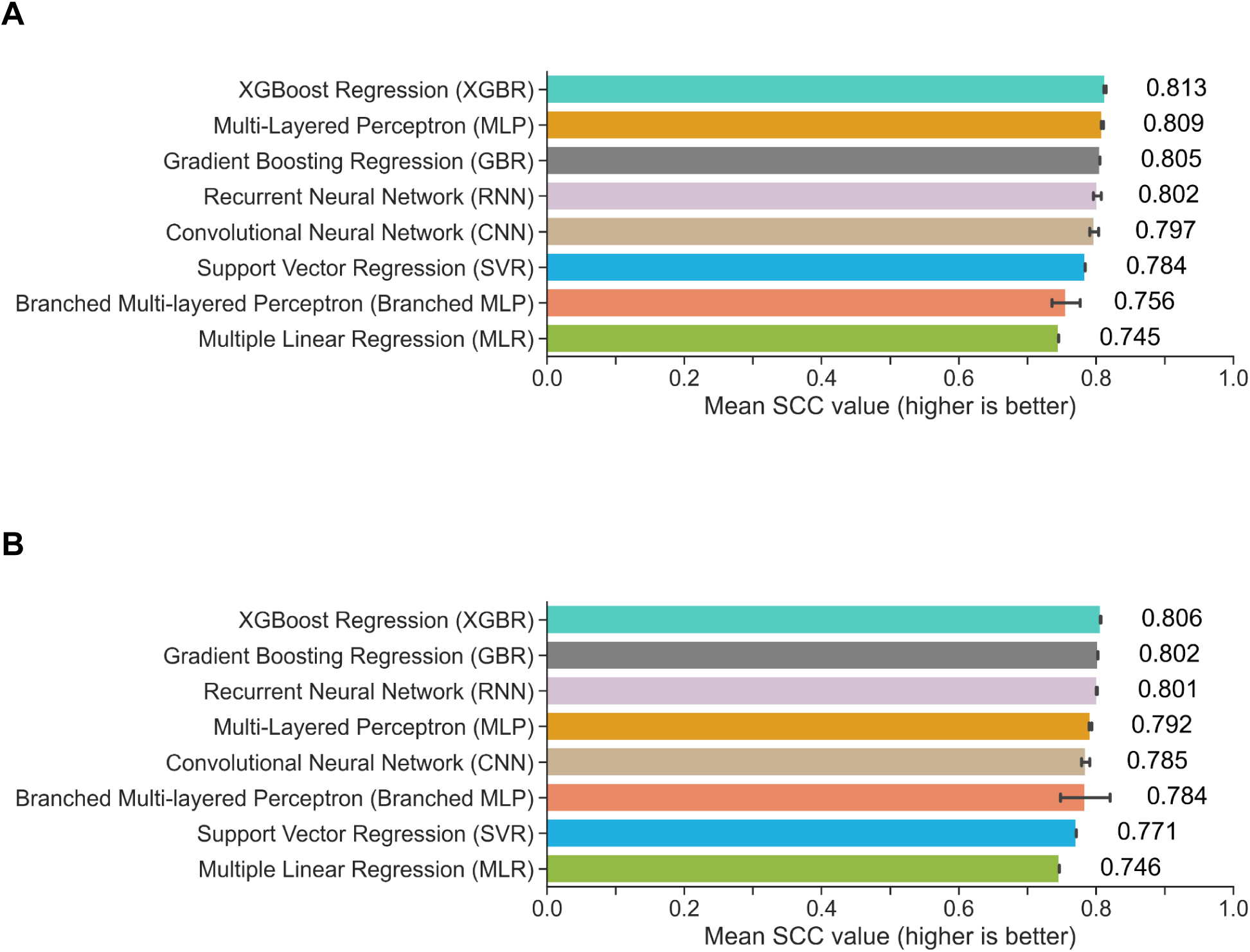
Spearman Correlation Coefficient experimental results. The SCC results depicted here support our PCC results for both prediction directions where we found our cross-patient XGBoost-based model maintained the highest performance.. A) The mean model performance values and their standard deviations (error bars) over 10 experimental runs are shown for GSC1 as the training and GSC2 as the testing dataset. B) The model SCC performance for the reverse prediction direction (GSC2 as testing and GSC1 as training).

### S6. Computational considerations

Script runtime was also taken into consideration as a measure of computational impact. The time it took for the experiments to be completed could provide some information towards its environmental impact and the availability of resources to make use of it. While we acknowledge the particular factors we needed to consider within our environment we also could not assume that future work in this space would not be influenced in some way by computational impact.

Although they were a factor, our computational times were not intended to be taken as absolute. These run times were measured according to the elapsed time for the python script containing the model and associated functions to run as calculated by the computational platform. For example, our deep learning scripts included functions for tasks such as gene level error recording, and visualization to name a few. Additionally, since the computational platform was a shared resource, run times can be influenced by the load of other laboratory tasks at a given time of day. Additionally, some of the model architectures make use of Graphical Processing Unit (GPU) acceleration which will impact many of the required calculations.

The model script runtime for this prediction direction found the MLR to complete the fastest followed by the average XGBR script completion time. The other models, with the exception of one, were comparatively close to both the MLR and XGBR times. Again, the Branched MLP performed behind the others. We provided information in the main body of this work as to how we went about recording runtimes. Additionally, we discussed how our data preprocessing was executed when the model scripts were run. Therefore, the Branched MLP is affected here by not only the fact that the 3-dimensional matrix of gene sequences were formed when the model was run but the one-hot encoding of that branch input happens as well. Together with the computational complexity, the runtime of the Branched MLP was over five times the MLR. It is of note that the XGBR model led the metric performance while the script completed the predictions faster than the majority of the others.

**Table S9.** Mean model script runtime measurements for experimental setup.

| model name | GSC1 (training) /<br>GSC2 (testing) ↓ | GSC2 (training) /<br>GSC1 (testing) ↓ |
| --- | --- | --- |
| XGBoost Regression (XGBR) | 7 min. 31 sec. | **7 min. 30 sec.** |
| Gradient Boosted Regression (GBR) | 10 min. 27 sec. | 7 min. 32 sec. |
| Multi-layered Perceptron (MLP) | 9 min. 12 sec. | 8 min. 38 sec. |
| Convolutional Neural Network (CNN) | 8 min. 58 sec. | 9 min. 49 sec. |
| Recurrent Neural Network (RNN) |  |  |
| Branched Multi-layered Perceptron (Branched MLP) | 43 min. 16 sec. | 38 min. 41 sec. |
| Support Vector Regression (SVR) | 8 min. 40 sec. | 9 min. 0 sec. |
| Multiple Linear Regression (MLR) | **7 min. 27 sec.** | 8 min. 1 sec. |

### S7. Supporting feature perturbation results

Our perturbation experiments included supplementary SCC metric calculations of the mean and standard deviations of the metric across the same set of experiments outlined in the main body of this work. As we observed from the analysis of our primary set of results, our SCC perturbation revealed a trend where the most pronounced effect to model performance occurred when the patients’ H3K27Ac feature values were perturbed. Just as with PCC, when we consider the prediction direction, perturbing this feature within GSC2 gave us the greatest reaction. The same feature for GSC1 and the opposite prediction direction gave us a smaller difference but still one that was more pronounced than the other features in that set. The differences being 76.208% against 15.879% (between the original and the perturbation results of 0.619646 and 0.128064 respectively). Interestingly, the perturbation of CTCF when GSC2 was the test set caused an increase in the metric by 0.000354. One explanation for this may be the fact that the models were not optimized for SCC.

Our takeaway here is that these results support our PCC results because we found H3K27Ac perturbation to cause the greatest performance change positively or negatively. The other features are impactful as well, albeit to lesser degrees.

**Fig S5A & S5B.**
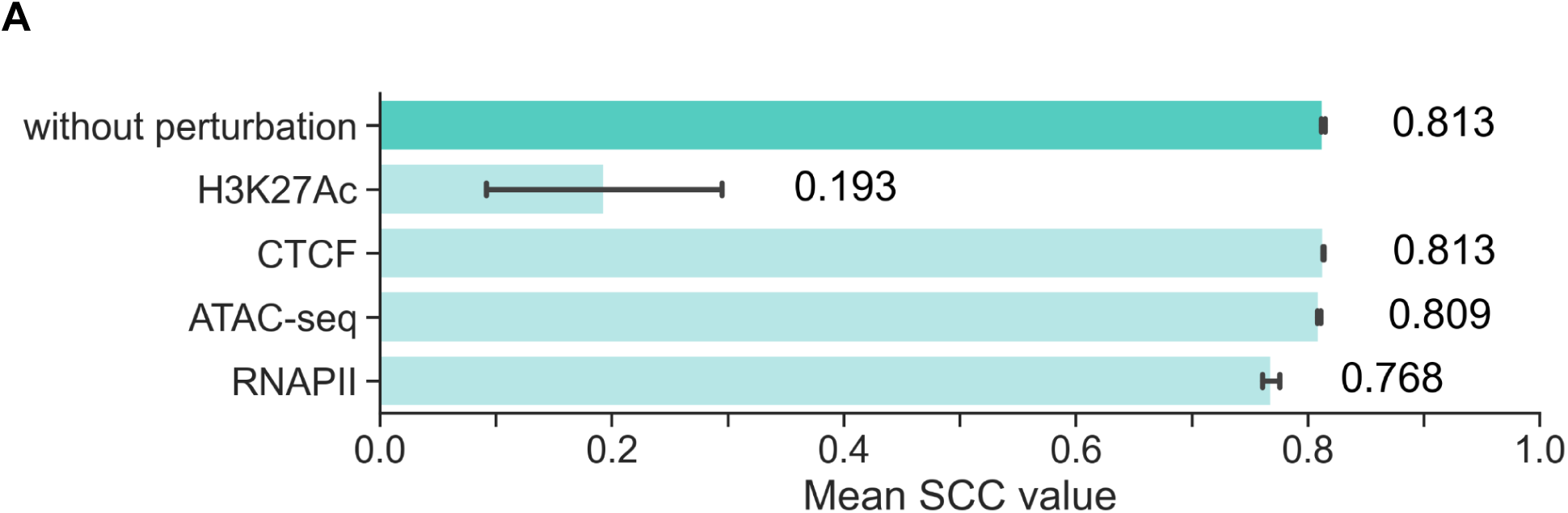
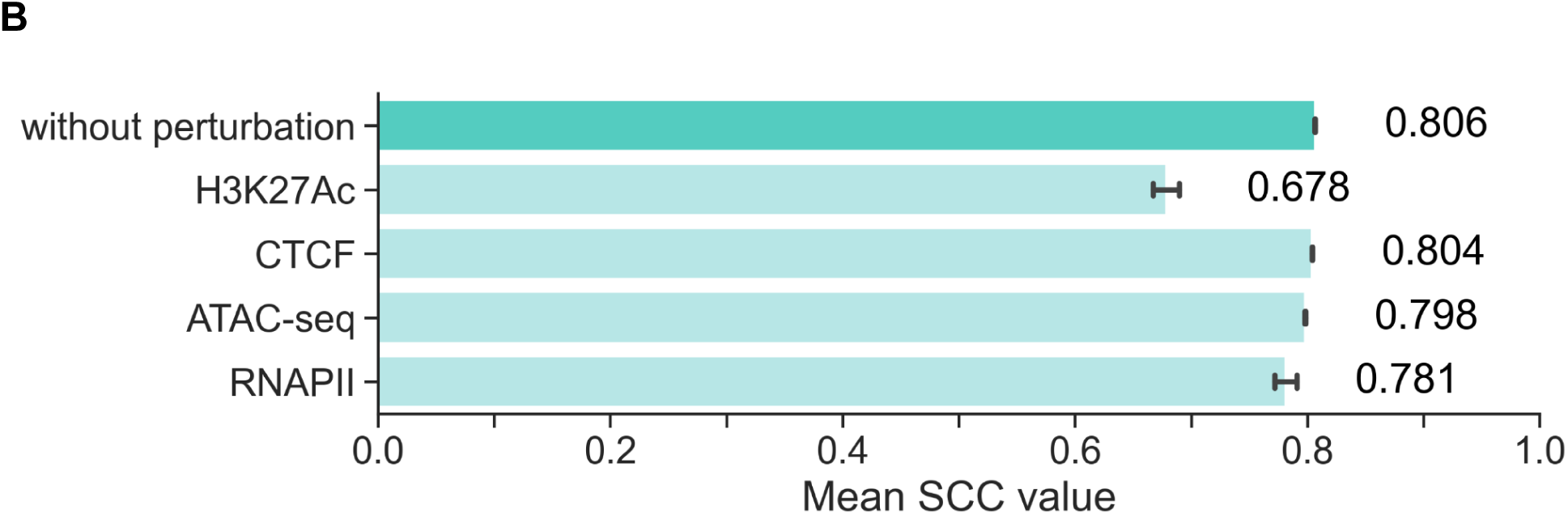
Spearman Correlation Coefficient results under feature perturbation. Our perturbation experiments produced SCC metric results alongside PCC results (Fig 4A and 4B). The results were compiled over the same 10 experimental runs, using our cross-patient XGBoost-based model and standard deviation indicated with error bars. A) Each epigenetic signal was perturbed separately when the model was trained with GSC1 and tested with GSC2. The model performance when H3K27Ac signals were perturbed was significantly decreased in comparison to the other signals. B) A decrease in performance was noted across all markers and particularly with H3K27Ac (GSC2 as training and GSC1 as testing).

### S8. Model feature importances support perturbation analysis

To investigate the interpretability of our model’s predictions we extracted feature importance calculations using the functionality incorporated into xgboost’s library. Since the datasets were flattened into a 2-dimensional form prior to model input, the model saw each gene’s bin values as the data’s features. When the model calculates feature importance, it does so for each bin. Therefore, we summed the individual feature importances corresponding to the four epigenetic markers (Fig S6A and S6B).

We found that the aggregated H3K27Ac marker was the most important feature for the model by considerable margins in both prediction directions. This supports our observations of the profound effects perturbation of the H3K27Ac signals had on model performance (Fig 5A and 5B, Fig S5A and S5B).

**Fig S6A & S6B.**
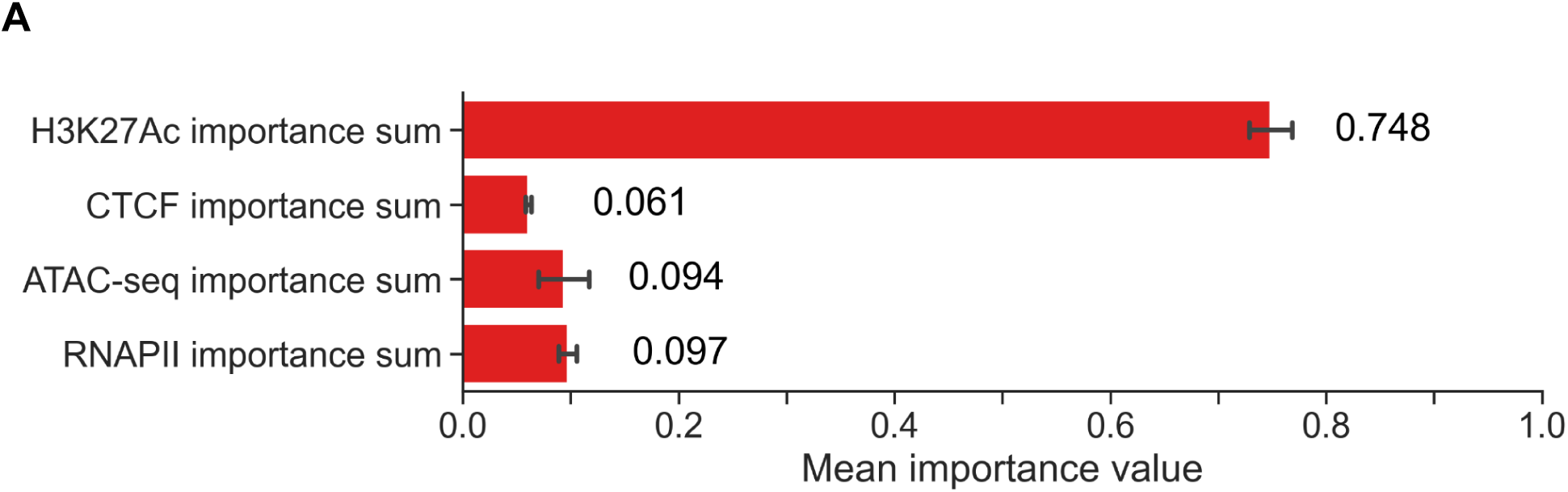
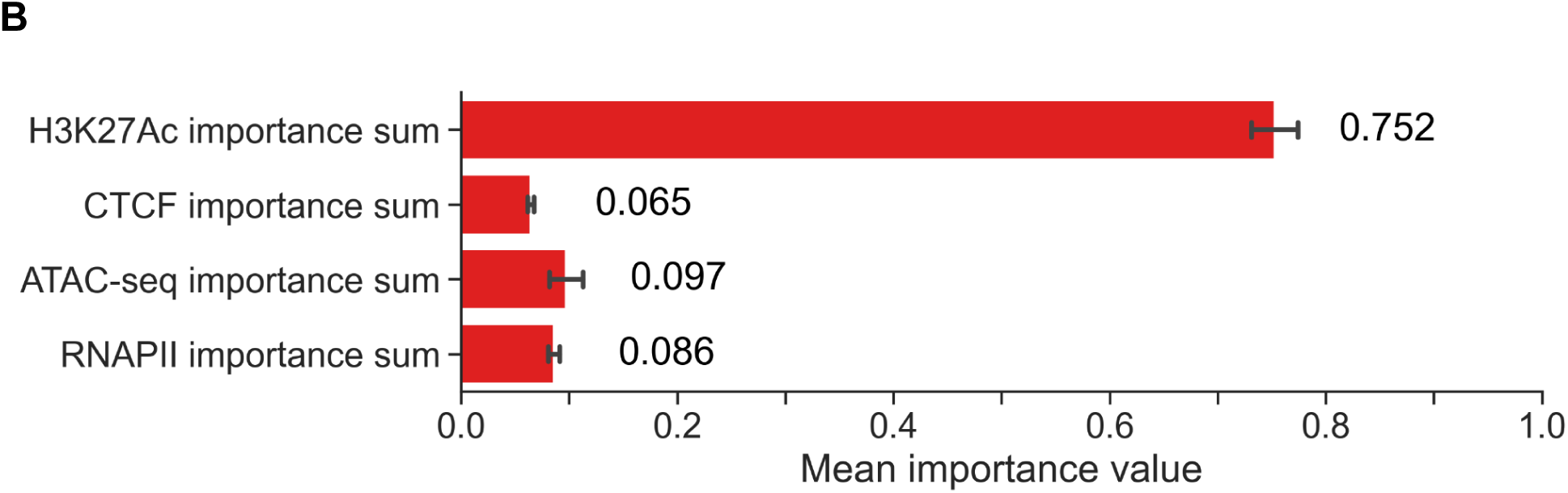
Feature importances extracted from our cross-patient xgboost-based model for GSC1→GSC2 (A) and GSC2→GSC1 (B). The model found the H3K27Ac features to be the most important for prediction of RNA-seq. The results visualized are the means over 10 experimental runs in each prediction direction with the error bars denoting the standard deviation.

### S9. How robust can our cross-patient prediction approach be when new patient data is introduced?

A guiding concept for our cross-patient computational approach was to uncover how well it would generalize to other glioblastoma patient data. The strength of our metric results alone answered that question. All of our cross-patient experiments were done with a test set derived from a different patient than the one used for that experiment’s tuning, training and validation. The hyperparameters chosen were kept consistent when we reversed the prediction direction and trained the model with the data of the second patient and evaluated it with that of the first. With this setup, metric performance of the “reversed” (GSC2 training/validation and GSC1 testing) prediction direction was comparable to the other results. We reasoned that this generalizability was observable given the fact that the training and testing of the model are using data from separate distributions. This is different from traditional machine learning experimental setups where the train, validation and test data are all derived from the same distribution and the datasets are created using cross-validation splitting techniques. This finding added support to our conjecture towards our method’s ability to generalize to unseen data.

From this observation, a question emerged of how sensitive was model performance to hyperparameter choices. We re-formulated the question and considered that since the hyperparameters were originally tuned when training and validation was done with GSC1’s data, would overall model performance be significantly affected if we instead hyperparameter tuned using GSC2’s data?

To answer this question we altered our experimental methodology. Instead of keeping the hyperparameters consistent for the “reversed” prediction direction we performed hyperparameter tuning using our XGBoost-based model. We did however, use the same values in the search set as the previous tuning for this model. This led us to choose hyperparameter values that were optimized again for PCC but were specific for this prediction direction. We then performed our new series of experiments with the same steps as our previous tests. When compared to our first set of tests with the same model and prediction direction, we found that there is a measurable increase in PCC and SCC by 0.002 in each.

To ensure that this observation was not due to the particular model of choice, we performed the same series of experiments using our MLP-based model. Again, we noted that when the hyperparameters for this specific prediction direction were chosen, there was an increase in the metrics (0.009 and 0.008 for PCC and SCC respectively).

Although there are minor performance advantages to hyperparameter tuning our models specifically for the direction of cross-patient prediction, the fact that the performance of the first experiments are within range of this optimized version is significant. Our analysis shows us that this approach when applied to our datasets, was generalizable and does not call the gene level predictions into question. We assert that these outcomes strengthen our understanding of the robustness of our cross-patient prediction methodology.

## Acknowledgments

For the neural crest and progenitor cell data, we acknowledge the ENCODE Consortium and Bradley Berstein lab, which the data originally came from. This research was conducted using computational resources and services at the Center for Computation and Visualization, Brown University. We are grateful to the members of COBRE-CBHD Computational Biology Core at Brown University and Eduardo Fajardo at Albert Einstein College of Medicine for the support. Y.S. was supported by Honjo International Foundation Fellowship. N.T. greatly acknowledge support from Warren Alpert Foundation. Effort for R.S and H.B was funded by the NIH award 1R35HG011939-01.

## References

1. Lathia JD, Mack SC, Mulkearns-Hubert EE, Valentim CL, Rich JN. Cancer stem cells in glioblastoma. Genes Dev. 2015 Jun 15;29(12):1203–17.

2. Patel AP, Tirosh I, Trombetta JJ, Shalek AK, Gillespie SM, Wakimoto H, et al. Single-cell RNA-seq highlights intratumoral heterogeneity in primary glioblastoma. Science. 20140612th ed. 2014 Jun 20;344(6190):1396–401.

3. Ignatova TN, Kukekov VG, Laywell ED, Suslov ON, Vrionis FD, Steindler DA. Human cortical glial tumors contain neural stem-like cells expressing astroglial and neuronal markers in vitro. Glia. 2002 Sep;39(3):193–206.

4. Liu G, Yuan X, Zeng Z, Tunici P, Ng H, Abdulkadir IR, et al. Analysis of gene expression and chemoresistance of CD133+ cancer stem cells in glioblastoma. Mol Cancer. 20061202nd ed. 2006 Dec 2;5(1):67.

5. Singh SK, Hawkins C, Clarke ID, Squire JA, Bayani J, Hide T, et al. Identification of human brain tumour initiating cells. Nature. 2004 Nov 18;432(7015):396–401.

6. Wang S, Zang C, Xiao T, Fan J, Mei S, Qin Q, et al. Modeling cis-regulation with a compendium of genome-wide histone H3K27ac profiles. Genome Res. 2016 Oct;26(10):1417–29.

7. Cheng C, Yan KK, Yip KY, Rozowsky J, Alexander R, Shou C, et al. A statistical framework for modeling gene expression using chromatin features and application to modENCODE datasets. Genome Biol. 2011;12(2):R15.

8. Schmidt F, Kern F, Schulz MH. Integrative prediction of gene expression with chromatin accessibility and conformation data. Epigenetics Chromatin. 2020 Feb 6;13(1):4.

9. Ouyang Z, Zhou Q, Wong WH. ChIP-Seq of transcription factors predicts absolute and differential gene expression in embryonic stem cells. Proc Natl Acad Sci U S A. 2009 Dec 22;106(51):21521–6.

10. Karlić R, Chung HR, Lasserre J, Vlahovicek K, Vingron M. Histone modification levels are predictive for gene expression. Proc Natl Acad Sci U S A. 2010 Feb 16;107(7):2926–31.

11. Kubo N, Ishii H, Xiong X, Bianco S, Meitinger F, Hu R, et al. Promoter-proximal CTCF binding promotes distal enhancer-dependent gene activation. Nat Struct Mol Biol. 2021 Feb;28(2):152–61.

12. Liu Y, Wu Z, Zhou J, Ramadurai DKA, Mortenson KL, Aguilera-Jimenez E, et al. A predominant enhancer co-amplified with the SOX2 oncogene is necessary and sufficient for its expression in squamous cancer. Nat Commun. 2021 Dec 8;12(1):7139.

13. Singh R, Lanchantin J, Sekhon A, Qi Y. Attend and Predict: Understanding Gene Regulation by Selective Attention on Chromatin. Adv Neural Inf Process Syst. 2017 Dec;30:6785–95.

14. Chen Y, Xie M, Wen J. Predicting gene expression from histone modifications with self-attention based neural networks and transfer learning. Front Genet. 2022;13:1081842.

15. Lee D, Yang J, Kim S. Learning the histone codes with large genomic windows and three-dimensional chromatin interactions using transformer. Nat Commun. 2022 Nov 5;13(1):6678.

16. Karbalayghareh A, Sahin M, Leslie CS. Chromatin interaction-aware gene regulatory modeling with graph attention networks. Genome Res. 2022 May;32(5):930–44.

17. Bigness J, Loinaz X, Patel S, Larschan E, Singh R. Integrating Long-Range Regulatory Interactions to Predict Gene Expression Using Graph Convolutional Networks. J Comput Biol J Comput Mol Cell Biol. 2022 May;29(5):409–24.

18. Massa AT, Mousel MR, Herndon MK, Herndon DR, Murdoch BM, White SN. Genome-Wide Histone Modifications and CTCF Enrichment Predict Gene Expression in Sheep Macrophages. Front Genet. 2020;11:612031.

19. Read DF, Cook K, Lu YY, Le Roch KG, Noble WS. Predicting gene expression in the human malaria parasite Plasmodium falciparum using histone modification, nucleosome positioning, and 3D localization features. PLoS Comput Biol. 2019 Sep;15(9):e1007329.

20. ENCODE Project Consortium. An integrated encyclopedia of DNA elements in the human genome. Nature. 2012 Sep 6;489(7414):57–74.

21. Luo Y, Hitz BC, Gabdank I, Hilton JA, Kagda MS, Lam B, et al. New developments on the Encyclopedia of DNA Elements (ENCODE) data portal. Nucleic Acids Res. 2020 Jan 8;48(D1):D882–9.

22. Hitz B, Kagda M, Lam B, Litton C, Small C, Sloan C, et al. Data navigation on the ENCODE Portal [Internet]. 2023 [cited 2024 Jun 23]. Available from: https://www.researchsquare.com/article/rs-3088639/v1

23. Hitz BC, Lee JW, Jolanki O, Kagda MS, Graham K, Sud P, et al. The ENCODE Uniform Analysis Pipelines.

24. Epigenome-based splicing prediction using a recurrent neural network-PMC [Internet]. [cited 2024 Jun 23]. Available from: https://www.ncbi.nlm.nih.gov/pmc/articles/PMC7343189/

25. Zhang J, Lee D, Dhiman V, Jiang P, Xu J, McGillivray P, et al. An integrative ENCODE resource for cancer genomics. Nat Commun. 2020 Jul 29;11(1):3696.

26. Meuleman W, Muratov A, Rynes E, Halow J, Lee K, Bates D, et al. Index and biological spectrum of human DNase I hypersensitive sites. Nature. 2020;584(7820):244–51.

27. Zwiener I, Frisch B, Binder H. Transforming RNA-Seq Data to Improve the Performance of Prognostic Gene Signatures. Emmert-Streib F, editor. PLoS ONE. 2014 Jan 8;9(1):e85150.

28. Kapoor S, Narayanan A. Leakage and the Reproducibility Crisis in ML-based Science [Internet]. arXiv; 2022 [cited 2023 Apr 9]. Available from: http://arxiv.org/abs/2207.07048

29. Mack SC, Singh I, Wang X, Hirsch R, Wu Q, Villagomez R, et al. Chromatin landscapes reveal developmentally encoded transcriptional states that define human glioblastoma. J Exp Med. 20190404th ed. 2019 May 6;216(5):1071–90.

30. Carén H, Stricker SH, Bulstrode H, Gagrica S, Johnstone E, Bartlett TE, et al. Glioblastoma Stem Cells Respond to Differentiation Cues but Fail to Undergo Commitment and Terminal Cell-Cycle Arrest. Stem Cell Rep. 2015 Nov 10;5(5):829–42.

31. Dirkse A, Golebiewska A, Buder T, Nazarov PV, Muller A, Poovathingal S, et al. Stem cell-associated heterogeneity in Glioblastoma results from intrinsic tumor plasticity shaped by the microenvironment. Nat Commun. 2019 Apr 16;10(1):1787.

32. Auffinger B, Tobias AL, Han Y, Lee G, Guo D, Dey M, et al. Conversion of differentiated cancer cells into cancer stem-like cells in a glioblastoma model after primary chemotherapy. Cell Death Differ. 2014 Jul;21(7):1119–31.

33. Grinsztajn L, Oyallon E, Varoquaux G. Why do tree-based models still outperform deep learning on tabular data? [Internet]. arXiv; 2022 [cited 2023 Mar 29]. Available from: http://arxiv.org/abs/2207.08815

34. Hu Y, Jiang Y, Behnan J, Ribeiro MM, Kalantzi C, Zhang MD, et al. Neural network learning defines glioblastoma features to be of neural crest perivascular or radial glia lineages. Sci Adv. 8(23):eabm6340.

35. Genome-wide analysis of polymerase III–transcribed Alu elements suggests cell-type–specific enhancer function-PMC [Internet]. [cited 2024 Jun 23]. Available from: https://www.ncbi.nlm.nih.gov/pmc/articles/PMC6724667/

36. Random Forests(TM) in XGBoost — xgboost 2.0.3 documentation [Internet]. [cited 2024 Jun 10]. Available from: https://xgboost.readthedocs.io/en/stable/tutorials/rf.html

37. XGBoost Parameters — xgboost 2.0.3 documentation [Internet]. [cited 2024 Jun 10]. Available from: https://xgboost.readthedocs.io/en/stable/parameter.html

